# WNT4 and WNT3A activate cell autonomous Wnt signaling independent of PORCN or secretion

**DOI:** 10.1101/333906

**Authors:** Deviyani M. Rao, Evelyn K. Bordeaux, Tomomi M. Yamamoto, Benjamin G. Bitler, Matthew J. Sikora

## Abstract

The enzyme PORCN is considered essential for Wnt secretion and signaling, however, we observed PORCN inhibition did not phenocopy the effects of WNT4 knockdown in WNT4-dependent breast cancer cells. This suggests a unique relationship between PORCN and WNT4 signaling. To examine the role of PORCN in WNT4 signaling, WNT4 or WNT3A were over-expressed in breast and ovarian cancer, and fibrosarcoma cell lines. Conditioned medium from these lines, and co-culture systems, were used to assess the dependence of Wnt secretion and activity on critical Wnt secretion proteins PORCN and WLS. We observed that WLS was universally required for Wnt secretion and paracrine signaling. In contrast, the dependence of WNT3A secretion and activity on PORCN varied across cell lines, and WNT4 secretion was PORCN-independent in all models. Surprisingly, WNT4 did not present paracrine activity in any tested context. Absent the expected paracrine activity of secreted WNT4, we identified cell autonomous Wnt signaling activation by WNT4 and WNT3A, independent of PORCN or secretion. The PORCN-independent, cell-autonomous Wnt signaling demonstrated herein may be critical in WNT4-driven cellular contexts, or those that are otherwise considered to have dysfunctional Wnt signaling.

**Summary Statement:** Wnt proteins can mediate an atypical mode of cell-autonomous signaling, distinct from paracrine signaling, that is independent of both palmitoylation by PORCN and Wnt secretion.

## Introduction

Wnt signaling is an ancestrally conserved pathway that plays fundamental roles in embryonic development and adult tissue homeostasis. Dysregulation of Wnt signaling is a causative factor for a range of human pathologies, including several forms of cancer (reviewed in (1)). As a result, inhibition of Wnt signaling has become an attractive therapeutic target in ongoing clinical trials, with some strategies targeting the upstream activation of signaling by Wnt proteins (1–3). Wnt proteins comprise a family of secreted glycoproteins that act as intercellular ligands, which stimulate a myriad of signal transduction cascades regulating cellular proliferation, stem cell renewal, cell motility, angiogenesis, and apoptosis (1, 4–6). Wnt proteins are post-translationally modified by the O-acyltransferase Porcupine (PORCN), which palmitoylates Wnt proteins at single serine residues (2, 7–9). This lipidation forms a binding motif for interaction with Wntless (WLS), which chaperones Wnt proteins to the plasma membrane for secretion (8, 10, 11). Once secreted, Wnt proteins signal in a paracrine manner, binding nearby receptor complexes. Wnts typically bind a Frizzled (FZD) receptor in conjunction with the LRP5 or LRP6 co-receptor, resulting in activation of the Disheveled second messenger proteins (DVL1/2/3 in humans) and initiation of either canonical (β-catenin-dependent) or non-canonical (β-catenin-independent) signaling (1, 4). The essential initiating step in Wnt processing is palmitoylation by PORCN, which has prompted the development of PORCN inhibitors, including IWP compounds (11), WNT974 (a.k.a. LGK974) (3), and others (2, 12). PORCN inhibitors have been shown to block Wnt secretion, inhibit downstream Wnt signaling, and suppress Wnt-driven tumor growth in animal models (3, 13, 14), with WNT974 currently in Phase I/II clinical trials for cancer treatment (NCT01351103, NCT02278133). Based on these observations, PORCN inhibitors are an attractive strategy to target Wnt-driven pathologies.

The Wnt protein WNT4 is critical in organogenesis of endocrine organs and regulation of bone mass, and underlies steroid hormone-related phenotypes in humans (15–22). WNT4 dysregulation via loss-of-function mutation results in developmental female to male sex reversal (23–26). Similarly, *WNT4* polymorphisms are associated with endocrine dysfunction, gynecological malignancies, reduced bone density with premature skeletal aging, and related phenotypes (27–33). WNT4 is also critical in mammary gland development, as *Wnt4* knockout in mouse mammary gland prevents progesterone-driven ductal elongation and branching during pregnancy (34, 35). In this context, activated progesterone receptor drives expression of *Wnt4* in mammary gland luminal cells resulting in paracrine signaling that supports maintenance of the mammary stem cell niche (6, 36–38). Despite these observed critical roles of WNT4 in both normal and malignant tissues, WNT4 signaling is crudely understood due to varied context-dependent functions. In a cell type- and tissue-specific manner, WNT4 (human or murine) has been shown to regulate either canonical or non-canonical Wnt signaling, and has been shown to either activate or suppress signaling (described in references herein). Further, conflicting reports exist as to whether Wnt4 can or cannot activate canonical Wnt signaling in the murine mammary gland (36, 39). As such, WNT4 has been described as a “problem child” among Wnt proteins. It is also unclear which FZD receptor complexes are utilized by WNT4, as WNT4 is often required for distinct, non-redundant functions versus other Wnt proteins (reviewed in (35)). Since WNT4 has myriad downstream signaling effects, inhibition of WNT4 upstream of Wnt effector pathways (e.g. with PORCN inhibitors) is an attractive approach to block WNT4 signaling in a “pathway indifferent” manner to treat WNT4-related pathologies.

We recently reported that regulation of *WNT4* expression is co-opted by the estrogen receptor in a subtype of breast cancer, invasive lobular carcinoma (ILC) (40, 41). Estrogen-driven WNT4 is required in ILC cells for estrogen-induced proliferation and survival, as well as anti-estrogen resistance (41). Though WNT4-driven signaling in ILC is yet to be fully elucidated, ILC cells lack the capacity to engage canonical Wnt signaling, as the characteristic genetic loss of E-cadherin in ILC leads to loss of β-catenin protein (41, 42). This suggests WNT4 drives non-canonical Wnt signaling in ILC cells. Though the specific non-canonical pathway activated by WNT4 is unknown, PORCN inhibition should be an effective strategy to block WNT4 upstream and treat this subtype of breast cancer. However, treatment of ILC cells with PORCN inhibitors did not suppress growth or survival. These unexpected results initiated further studies into the mechanisms of WNT4 secretion and signaling. In this report, we show WNT4 secretion is mediated by atypical mechanisms. Our observations challenge the paradigm that PORCN-mediated secretion is required for Wnt signaling, and suggest a novel process by which Wnt proteins, including WNT4, can initiate non-canonical Wnt signaling.

## Results

### PORCN inhibition does not mimic WNT4 siRNA in lobular carcinoma cells

We hypothesized that since ILC cells are dependent on WNT4 for proliferation and survival (41), inhibition of PORCN would phenocopy *WNT4* siRNA by blocking WNT4 secretion and downstream signaling. Proliferation and cell death were monitored by live cell imaging of MM134 (ILC) cells either transfected with siRNA targeting *PORCN* (siPORCN) or treated with PORCN inhibitor (PORCNi) LGK974. Proliferation and cell death were compared to untreated cells, and cells treated with the antiestrogen fulvestrant (Fulv) or transfected with siRNA targeting *WNT4* (siWNT4). As we previously reported (41), siRNA-mediated *WNT4* knockdown or Fulv halt proliferation, and WNT4 knockdown induces cell death (**Fig. 1A**). However, neither genetic nor chemical PORCN inhibition had any effect on cell proliferation or survival of MM134 cells (**Fig. 1A,B**). Similar results were obtained in ILC cell line SUM44PE, as PORCN inhibitor at concentrations up to 1μM did not affect proliferation (**Supplemental Fig. 1**). These data suggest PORCN inhibition is not sufficient to inhibit WNT4 function, and WNT4 signaling likely occurs via PORCN-independent mechanisms.

**Figure 1.**
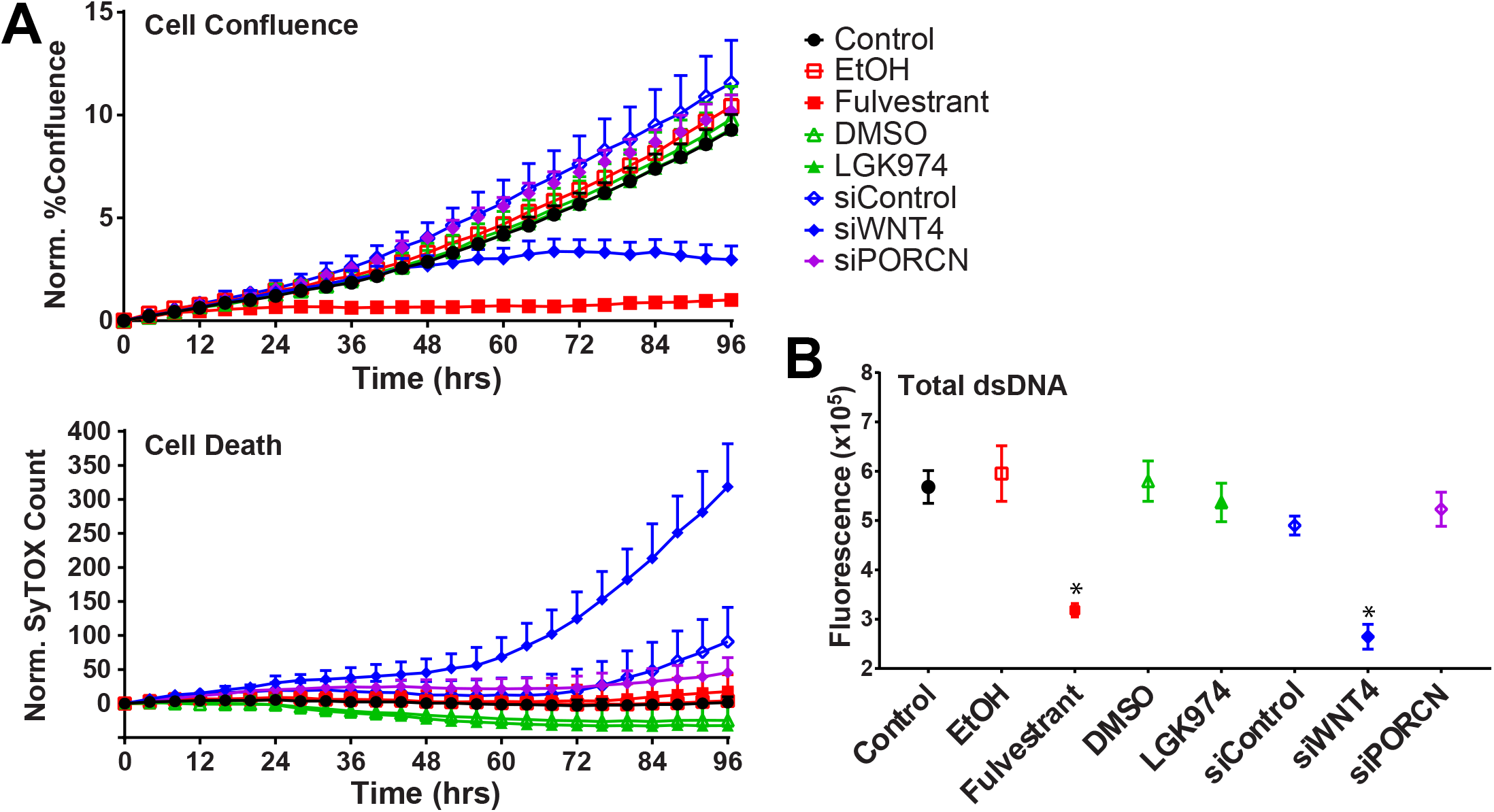
Inhibition or knockdown of PORCN does not phenocopy knockdown of WNT4 in MM134. (A), MM134 cells were transfected with siRNA or treated with fulvestrant (100nM), LGK974 (10nM), or 0.1% vehicle (EtOH or DMSO) at time 0, prior to live cell imaging for proliferation (phase-contrast confluence) and death (SyTOX green incorporation). Points represent mean of 6 biological replicates ±SD. (B), Total double-stranded DNA was measured from assays in (A) at timecourse completion. *, p<0.05 vs control, ANOVA with Dunnett’s multiple correction. Results in A,B are representative of three independent experiments.

### WNT4 secretion is WLS-dependent but PORCN-independent

Since targeting PORCN did not phenocopy WNT4 knockdown, we further examined the role of PORCN in WNT4 secretion. To facilitate Wnt secretion studies we over-expressed WNT3A or WNT4 in MM134 (MM134:W3 and MM134:W4; **Fig. 2A, Table 1**), and measured secreted Wnt proteins in conditioned medium. Of note, since epitope tags may alter Wnt secretion and activity (e.g. (8)), we performed all studies with non-tagged Wnt constructs. A general workflow for experiments assessing Wnt secretion and function, for each figure, is shown in **Supplemental Fig. 2**.

**Figure 2.**
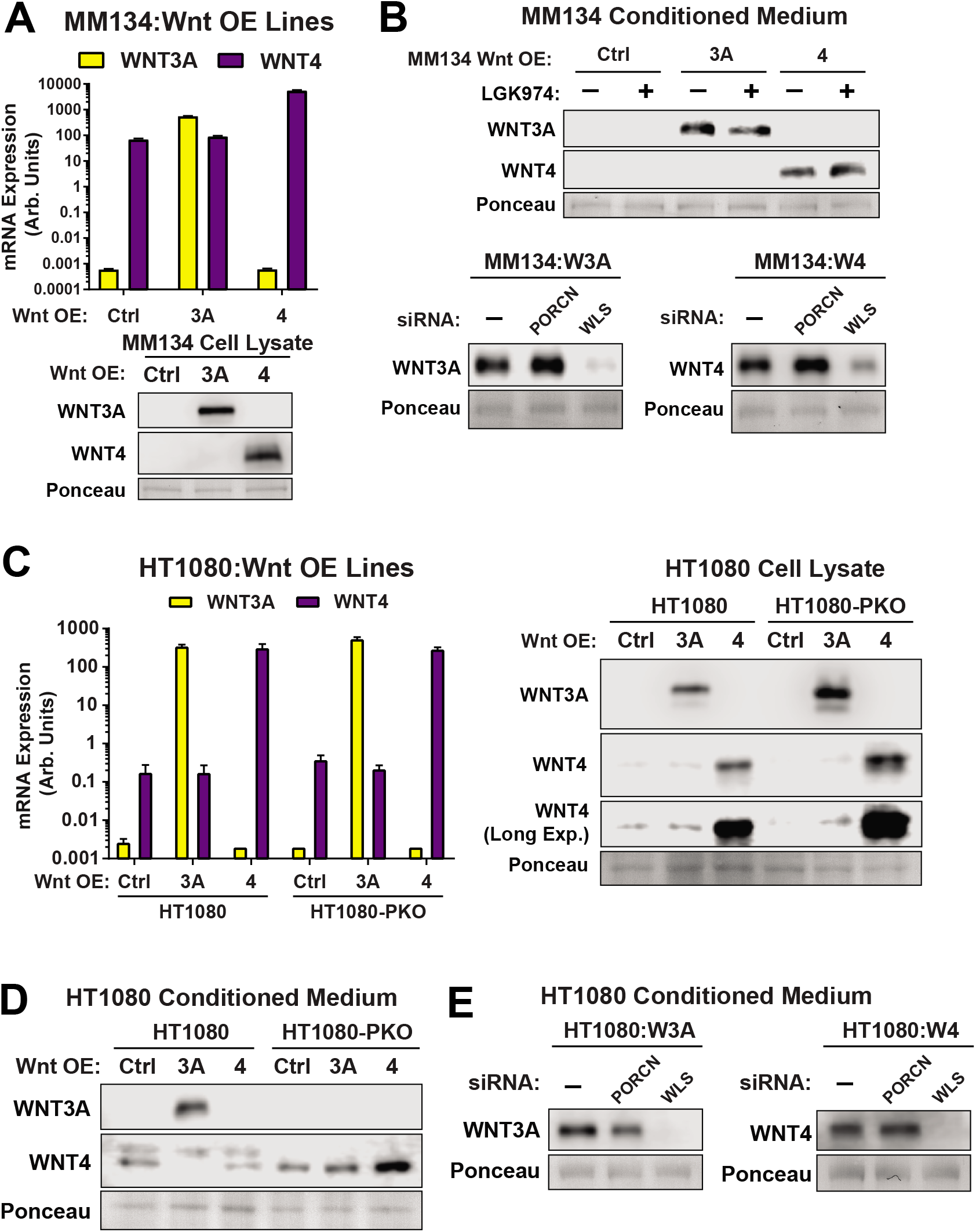
WNT4 secretion is PORCN-independent, but WLS-dependent. (A), MM134 constitutive Wnt over-expression cell lines were created as detailed in Methods and Materials. Top, qPCR for WNT3A and WNT4. Bars represent mean of 2 technical replicates ±SD. Bottom, immunoblot for cellular expression of WNT3A and WNT4. Endogenous WNT4 could not be visualized here due to the level of over-expression (B), Top, MM134 were treated with 10nM LGK974, and medium was allowed to condition for 7 days. Bottom, MM134 were transfected with siPORCN or siWLS, and after 24hrs, medium was changed and conditioned for 7 days. Total protein was extracted from medium as above for immunoblot. (C), HT1080 constitutive Wnt over-expression cell lines were created as detailed in Methods and Materials. Left, qPCR for WNT3A and WNT4. Bars represent mean of 3 technical replicates ±SD. Right, immunoblot for cellular expression of WNT3A and WNT4. Over-expression shows endogenous WNT4 expression. (D), HT1080 medium was conditioned for 5 days as described in Materials and Methods, prior to total protein extraction for immunoblot. (E), HT1080 cells were transfected with siPORCN or siWLS, and after 24hrs, medium was changed and conditioned for 5 days. Total protein was extracted from medium as above for immunoblot.

**Table 1 –.**
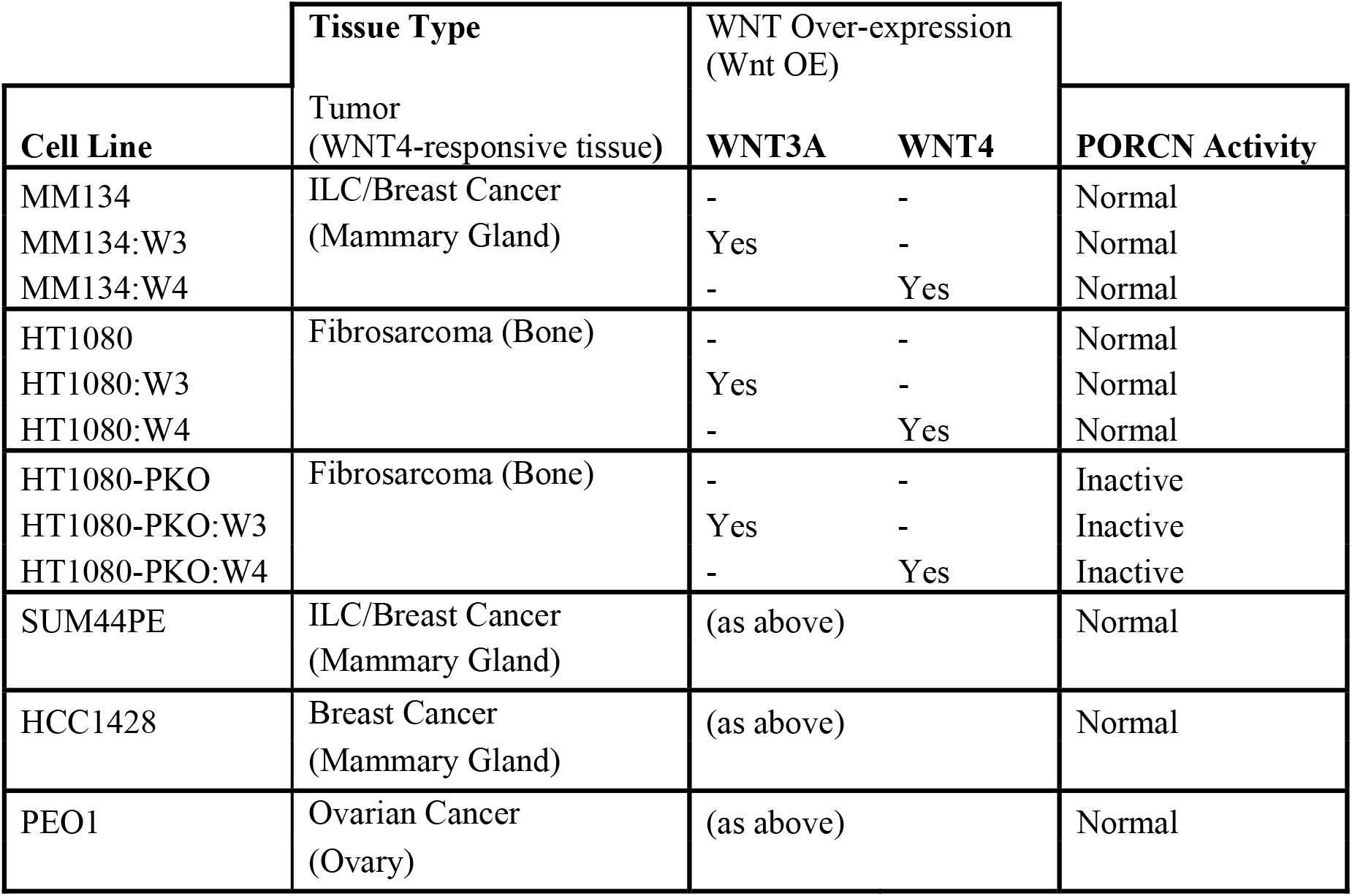
Model Systems

| Cell Line | Tissue Type<br>(WNT4-responsive tissue) | WNT Over-expression<br>(Wnt OE) |  | PORCN Activity |
| --- | --- | --- | --- | --- |
|  |  | WNT3A | WNT4 |  |
| MM134 | ILC/Breast Cancer<br>(Mammary Gland) | - | - | Normal |
| MM134:W3 |  | Yes | - | Normal |
| MM134:W4 |  | - | Yes | Normal |
| HT1080 | Fibrosarcoma (Bone) | - | - | Normal |
| HT1080:W3 |  | Yes | - | Normal |
| HT1080:W4 |  | - | Yes | Normal |
| HT1080-PKO | Fibrosarcoma (Bone) | - | - | Inactive |
| HT1080-PKO:W3 |  | Yes | - | Inactive |
| HT1080-PKO:W4 |  | - | Yes | Inactive |
| SUM44PE | ILC/Breast Cancer<br>(Mammary Gland) | (as above) |  | Normal |
| HCC1428 | Breast Cancer<br>(Mammary Gland) | (as above) |  | Normal |
| PEO1 | Ovarian Cancer<br>(Ovary) | (as above) |  | Normal |

PORCN-mediated palmitoylation of Wnt proteins is commonly described as required for Wnt binding to WLS and transport to the cell surface for secretion (see Introduction), so we examined the requisite of PORCN (using PORCNi and siPORCN) or WLS (using siWLS) for Wnt secretion. Secreted WNT3A and WNT4 were detected in conditioned medium from MM134:W3 and MM134:W4 respectively (**Fig. 2B**). Consistent with the lack of effect of cell proliferation, PORCNi treatment had no effect on WNT4 secretion, and WNT3A secretion was also unaffected by PORCNi (**Fig. 2B, top**). Similarly, siPORCN had no effect on secretion of either WNT4 or WNT3A (**Fig. 2B, bottom**; efficacy of PORCNi and siPORCN were confirmed below). However, WLS was required for Wnt secretion, as siWLS suppressed secretion of both WNT3A and WNT4 from MM134 (**Fig. 2B, bottom**). These data suggest that Wnt processing and secretion may be atypical in ILC cells, but the PORCN-independent secretion of WNT4 is a potential mechanism of PORCNi resistance (**Figure 1**).

To determine whether PORCN-independent WNT4 secretion is ILC-specific, we utilized the HT1080 fibrosarcoma cell line, a well-characterized model for Wnt secretion, signaling, and activity (8, 43). HT1080 are derived from a bone-like tissue and thus are a relevant context for WNT4 signaling (18, 22) (e.g. WNT4 activates DVL via non-canonical Wnt signaling in HT1080 (8)). We generated WNT3A and WNT4 over-expressing cells from both wild-type HT1080 and PORCN-knockout HT1080 (HT1080-PKO, clone delta-19 (43)) (**Fig. 2C, Table 1**), and assessed Wnt secretion as above. Unlike the ILC model, WNT3A secretion from HT1080 was PORCN-dependent, as WNT3A could be detected in conditioned medium from HT1080:W3 but not HT1080-PKO:W3 cells (**Fig. 2D**). PORCNi treatment also blocked WNT3A secretion from HT1080:W3 (**Supplemental Fig. 3A**). In contrast, WNT4 secretion was detected from both HT1080:W4 and HT1080-PKO:W4 (**Fig. 2D**), and PORCNi did not suppress WNT4 secretion from HT1080:W4 (**Supplemental Fig. 3A**), supporting that WNT4 secretion is PORCN-independent.

Endogenously expressed WNT4 was also detected in conditioned medium from HT1080 and HT1080-PKO cell lines (**Fig. 2D**). We observed that secreted WNT4 can be resolved by electrophoresis as a doublet. The larger species was PORCN-dependent and not detected in HT1080-PKO, suggesting that these species represent palmitoylated versus non-palmitoylated proteins. Notably, in HT1080:W3 endogenous secreted WNT4 shifted to the larger species (**Fig. 2D**, column 2), potentially due to positive feedback activation of PORCN activity (44). We observed loss of both species in conditioned media after *WNT4* knockdown by siRNA, confirming both secreted species as WNT4 (**Supplemental Fig. 3A**).

These data indicate that while WNT4 is modified by PORCN, PORCN is not required for WNT4 secretion. WLS knockdown suppressed secretion of both WNT3A and WNT4 from HT1080 (**Fig. 2E**), confirming that WNT3A secretion is dependent on both WLS and PORCN, while WNT4 secretion is WLS-dependent but PORCN-independent.

Though WNT4 secretion was PORCN-independent in both MM134 and HT1080, WNT4 appeared to be post-translationally modified during secretion (**Fig. 2D**). Also, in both MM134 and HT1080, secreted WNT4 migrated as a higher molecular weight species than WNT4 from cell lysate (**Supplemental Fig. 3B**). The increased molecular weight is due at least in part to glycosylation (45), as treatment with tunicamycin (N-linked glycosylation inhibitor) decreased the apparent molecular weight of secreted WNT4 in either cell line (**Supplemental Fig. 3B**). Notably, MM134 secreted two WNT4 species when treated with tunicamycin, suggesting WNT4 modification can be variable and cell context-specific.

Together these data demonstrate that WNT4 secretion is PORCN-independent, and that Wnt protein processing and signaling are more broadly atypical in ILC. PORCN-independent WNT4 secretion may explain the disparate effects of siWNT4 versus PORCN inhibition. However, it is unclear if Wnt proteins secreted independently of PORCN are competent to activate paracrine Wnt signaling.

### WNT4 over-expression does not activate paracrine Wnt signaling

Since palmitoylation of Wnt proteins is typically necessary for activation of downstream signaling, we assessed whether WNT4 secreted independent of PORCN could activate paracrine Wnt signaling. Paracrine signaling was tested by treating HT1080-PKO cells (“receiver” cells, PKO reduces endogenous paracrine Wnt signaling; see **Supplemental Fig. 2**) with conditioned medium from MM134 cells, with or without Wnt over-expression. After 24hr treatment with conditioned medium, activation of Wnt signaling in receiver cells was assessed via phosphorylation of DVL2, DVL3, and LRP6, and expression of β-catenin target gene *AXIN2.* Conditioned medium from neither MM134:W4 nor MM134:W3 were able to activate Wnt signaling in receiver cells by any measure (**Fig. 3A,B**) despite the presence of Wnt protein in conditioned medium from these cells (**Fig. 2B**). This suggests Wnt secretion is not sufficient for paracrine activity. We also assessed Wnt paracrine activity from wild-type vs PKO Wnt-overexpressing HT1080. Conditioned medium from HT1080:W3 activated Wnt signaling in the receiver cells by all measures (**Fig. 3C,D**), and this was blocked by PORCN knockout (ie. from HT1080-PKO:W3), consistent with a lack of WNT3A secretion in HT1080-PKO (**Fig. 2D**). However, conditioned medium from neither HT1080:W4 nor HT1080-PKO:W4 activated Wnt signaling in the receiver cells (**Fig. 3C,D**) despite detectable secreted WNT4 in both contexts (**Fig. 2D**). Thus, secreted WNT4 from MM134, HT1080, or HT1080-PKO was unable to activate paracrine Wnt signaling.

**Figure 3.**
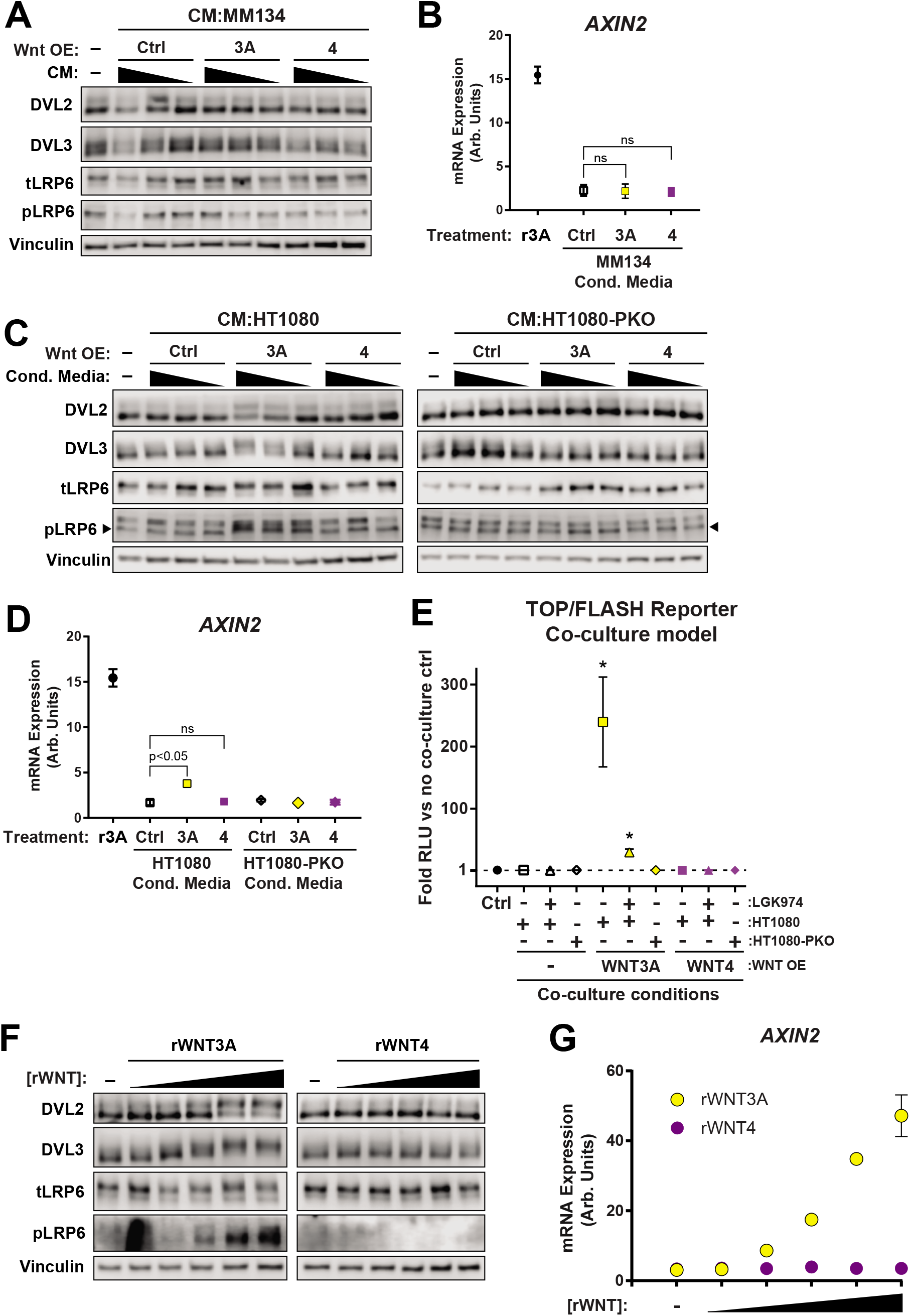
WNT4 does not activate paracrine Wnt signaling. (A-D) HT1080-PKO control cells were treated for 24 hours with conditioned media (CM, at 50%, 25% or 12.5% final volume supplemented with fresh medium) from either MM134 (A-B) or HT1080 cell lines (C-D). (A,C) Immunoblots of whole cell lysates from the treated HT1080-PKO cells were run and probed for DVL2, DVL3, total LRP6 and phosphorylated LRP6. (B,D) mRNA from the treated HT1080-PKO cells was extracted for qPCR for AXIN2 mRNA expression levels, vs RPLP0. Points represent mean of 2 technical replicates ±SD. (B,D) Cells treated with 62.5ng/mL rWNT3A as a positive control. Statistics obtained using ANOVA with Dunnett’s multiple correction. (E) HT1080-PKO cells were transfected with a TOP-FLASH reporter plasmid, then co-cultured with either HT1080 or HT1080-PKO WNT overexpressing (OE) cells, with or without 10nM LGK974. ‘Ctrl’ represents TOP-FLASH transfected HT1080-PKO without co-culture. WNT signaling activity, as measured by luminescence, was performed using a dual luciferase assay. Statistics obtained using Student’s unpaired t-test compared to the no co-culture control (ctrl). * represents p<0.005. Points represent mean of 3 technical replicates ±SD. Results are representative of two independent experiments. (F-G) HT1080-PKO control cells were treated for 24 hours with recombinant WNT protein (at concentrations of 10ng/ml, 50ng/ml, 100ng/ml, 250ng/ml, or 500ng/ml). (F) Immunoblots of whole cell lysates were run and (G) mRNA extracted for qPCR were performed as above (A-D). Statistics obtained using ANOVA with Dunnett’s multiple correction. Points represent mean of 2 technical replicates ±SD.

Paracrine signaling by Wnt proteins can be mediated by Wnt secretion as well as by Wnt presentation on the cell surface and subsequent cell-cell contact (46, 47). To determine whether paracrine WNT4 signaling is initiated specifically by cell-cell contact, we assessed whether Wnt signaling could be activated in “receiver” cells via co-culture with Wnt-overexpressing cells. HT1080-PKO cells transfected with the TOP-FLASH reporter (“receiver” cells) were co-cultured with cells expressing WNT3A or WNT4 (**Fig. 3E**). WNT3A-expressing cells activated TOP-FLASH in co-cultured cells in a PORCN-dependent manner (ie. blocked by LGK974, and absent from HT1080-PKO:W3 cells). However, under no condition was WNT4 able to activate TOP-FLASH in co-cultured cells. While WNT3A is able to mediate paracrine Wnt signaling in both secreted and co-culture models, neither secreted nor cell surface WNT4 mediates paracrine signaling.

Since WNT4 from neither HT1080 nor MM134 cells was able to activate paracrine Wnt signaling, we treated HT1080-PKO cells with recombinant human Wnt proteins (rWNT3A and rWNT4; 10-500ng/mL). rWNT3A activated Wnt signaling by all measures, while rWNT4 failed to activate Wnt signaling at any concentration (**Fig. 3F,G**). These data suggest that HT1080 may be non-responsive to paracrine WNT4, raising the need for orthogonal systems to validate the function of both secreted and recombinant WNT4. To address this, we used MC3T3-E1 as an additional “receiver” cell line (**Supplemental Fig. 2**). MC3T3-E1 cells are another bone-like model highly responsive to exogenous Wnt protein, and induce alkaline phosphatase production upon Wnt signaling activation (measured by colorimetric assay, see Materials and Methods (48)). Conditioned medium as above was used to treat “receiver” 3T3-E1 cells, and alkaline phosphatase (AP) activity was used as the readout for activation of paracrine Wnt signaling. Either rWNT3A or rWNT4 increased AP activity (**Fig. 4A**), and both rWNT3A and rWNT4 induced DVL and LRP6 phosphorylation in 3T3-E1 cells (**Supplemental Fig. 4A**). This confirmed that 3T3-E1 respond to paracrine WNT4, and could be used to assess the activity of secreted Wnts in conditioned medium. Conditioned medium from HT1080:W3, but not HT1080-PKO:W3, induced AP activity in 3T3-E1 (**Fig. 4B**), consistent with a requirement of PORCN for paracrine WNT3A signaling. However, though rWNT4 induced AP activity, conditioned medium from neither HT1080:W4 nor HT1080-PKO:W4 induced AP activity in 3T3-E1 (**Fig. 4B**). Parallel results were obtained using MM134, as conditioned medium from MM134:W3 induced AP activity in 3T3-E1, which was blocked by PORCNi treatment, but no AP activity was induced with MM134:W4 conditioned medium (**Fig. 4C**). We considered that the concentration of secreted WNT4 protein may be insufficient to activate 3T3-E1, but secreted WNT3A or WNT4 is within the range of active rWNT concentrations in this assay (**Supplemental Figure 4B-D**). Notably, commercially available rWNT3A and rWNT4 are produced lacking the N-terminal signal peptide, and both rWNT3A and rWNT4 migrated as a smaller peptide than the corresponding Wnt protein secreted from either HT1080 or MM134 (**Supplemental Fig. 4B-C**). Taken together, these results indicate secreted Wnt proteins are not equivalent to recombinant proteins, and have distinct capacities for activating paracrine Wnt signaling. However, while secreted WNT3A activated paracrine Wnt signaling in a context-dependent manner (based on both source and receiver cells), secreted WNT4 was unable to act as a functional paracrine signaling ligand in any tested context.

**Figure 4.**
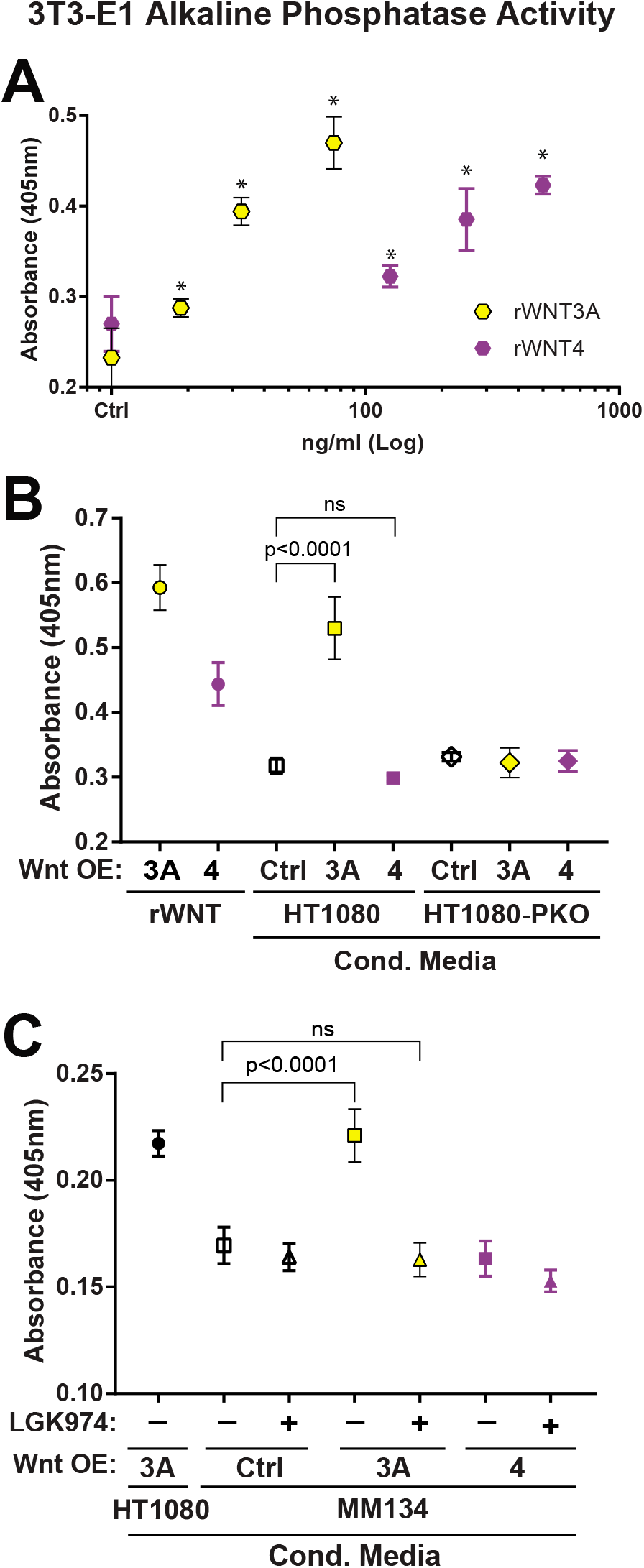
Secreted versus recombinant Wnt protein differentially activate paracrine signaling. (A-C), MC3T3-E1 cells were plated into a 96-well plate and 24hrs later were treated with (A), rWNT3A (0, 18.75, 32.5, 75ng/ml) or rWNT4 (0, 125, 250, 500ng/ml), (B), rWNT3A (62.5ng/ml), rWNT4 (250ng/ml) or 50% conditioned media (CM) from HT1080 cell lines, or (C), 50% CM from HT1080 or MM134 cell lines that were treated with or without 10nM LGK974. Points represent a mean of 4 (A), 3 (B) and 6 (C) biological replicates ±SD. Estimated concentrations of secreted WNTs (sWNT) present in CM based on linear regression of rWNTs is shown in Supplemental Fig. 4. *, p<0.05 vs control, ANOVA with Dunnett’s multiple correction.

### PORCN-independent WNT4 secretion and lack of paracrine activity is observed in varied model systems

We further examined Wnt secretion and function via WNT3A or WNT4 over-expression in a second ILC cell line (SUM44PE), an additional breast cancer cell line (HCC1428), and an ovarian cancer cell line (PEO1) (**Table 1**). Like bone, ovarian cancer is a relevant context as WNT4 mediates Müllerian tissue and ovary development (23, 25). Wnt expression and secretion were assessed as above, and paracrine activity of secreted Wnt proteins was tested using the 3T3-E1 receiver model.

Secreted WNT3A and WNT4 were detected after over-expression in SUM44PE (ILC, **Fig. 5A**), and WNT4 secretion in SUM44PE was PORCN-independent but WLS-dependent (**Fig. 5B**). Neither WNT3A nor WNT4 induced AP activity versus parental SUM44PE cells (**Fig. 5C**). These data with SUM44PE are consistent with our observations of PORCN-independent Wnt secretion and inactive paracrine activity of secreted Wnt proteins in ILC. Secreted WNT4 also lacked any paracrine activity from both HCC1428 (**Fig. 5D-E**) and PEO1 (**Fig. 5F-G**). In total, we are unable to detect paracrine WNT4 activity in 5 cell lines derived from 3 different WNT4-responsive tissues of origin (mammary gland, bone, and ovary), further supporting that secreted WNT4 does not mediate paracrine signaling in WNT4 expressing cells.

**Figure 5.**
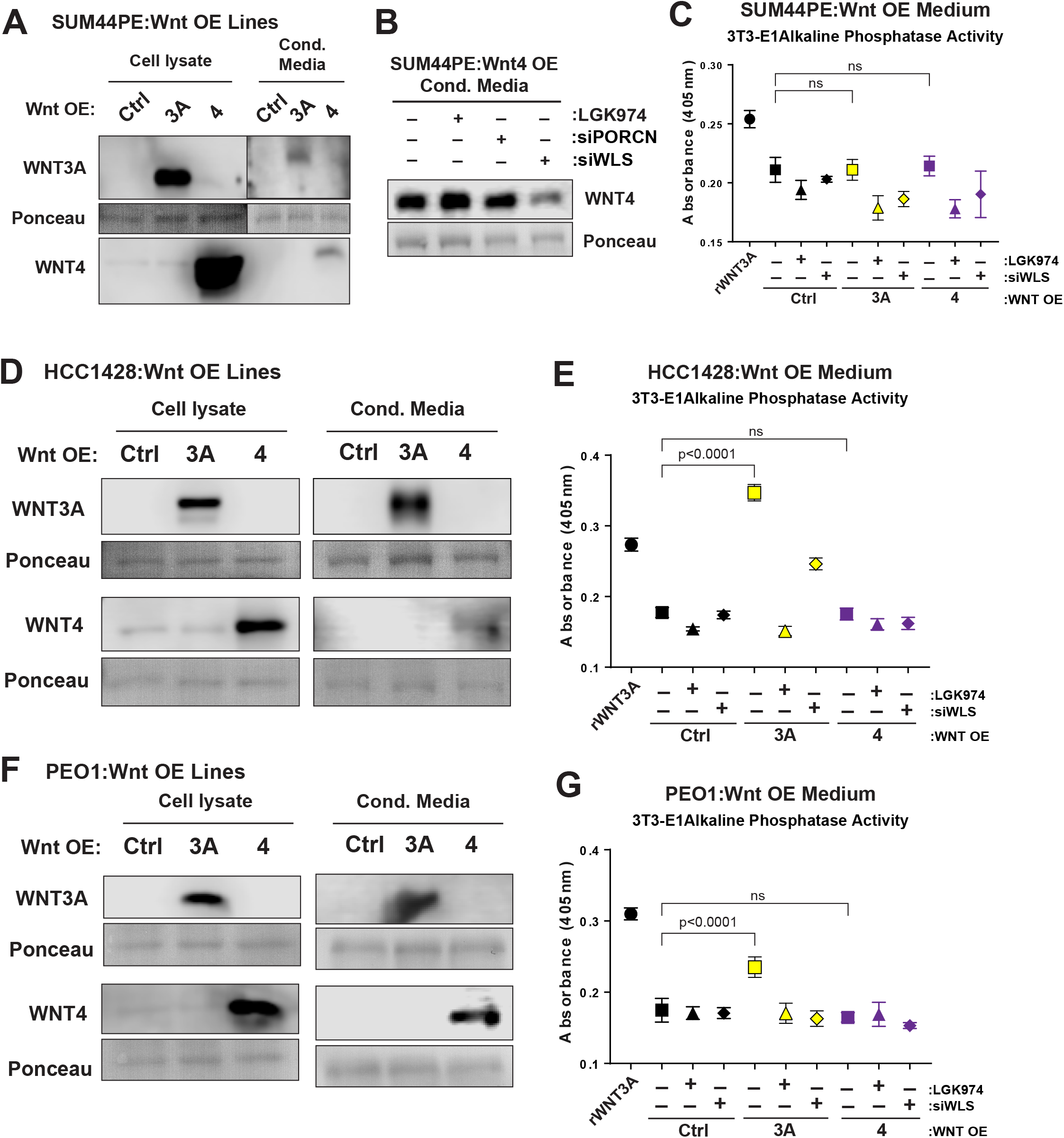
WNT4 from various cell types lacks paracrine activity. Western blots of whole cell lysate or conditioned media from WNT3A or WNT4 overexpressing (A) SUM44PE, (D) HCC1428, or (F) PEO1 cell lines were probed for either WNT3A or WNT4. (B), A western blot of conditioned media from SUM44PE WNT4 over expressing cells treated with either LGK974, siPORCN or siWLS. Statistics obtained using ANOVA with Dunnett’s multiple correction. Alkaline phosphatase production induced by conditioned media from WNT3A or WNT4 overexpressing (C) SUM44PE, (E) HCC1428 or (G) PEO1 cell lines was measured. Points represent a mean of 4 biological replicates ±SD.

### Secreted or paracrine WNT4 are not required for ILC cell proliferation and viability

Since secreted WNT4 did not activate paracrine Wnt signaling in MM134 or SUM44PE cells, we hypothesized that WNT4 secretion is dispensable for WNT4 function in ILC cells. To confirm this, we examined whether blocking WNT4 secretion by WLS knockdown would phenocopy WNT4 knockdown in MM134 cells. MM134 were transfected with siRNAs targeting WNT4, PORCN, or WLS, followed by cell proliferation and death assays as above. Despite the loss of WNT4 secretion, siWLS had no detrimental effect on MM134 proliferation or viability and did not phenocopy siWNT4 (**Supplemental Fig. 5A-B**). Similarly, we attempted to rescue WNT4 knockdown with conditioned medium from MM134 (with or without Wnt over-expression) or rWNT protein. Exogenous Wnt protein had no effect on siWNT4-mediated growth inhibition or cell death (**Supplemental Fig. 5C-D**). These data support that secretion may does not mediate the activation of WNT4-driven signaling.

### WNT4 and WNT3A activate cell-autonomous signaling

Our observations suggest WNT4 may activate signaling not via paracrine or autocrine mechanisms, but rather by a cell-autonomous mechanism. To examine cell-autonomous Wnt-induced signaling we assessed DVL and LRP6 phosphorylation and *AXIN2* expression directly in HT1080 and HT1080-PKO Wnt over-expressing cells (**Fig. S2; Fig. 6A,B**). Consistent with conditioned medium/receiver cell experiments, both HT1080:W3 and HT1080:W4 had increased DVL and LRP6 phosphorylation (**Fig. 6A**), but only HT1080:W3 had increased *AXIN2* expression (**Fig. 6B**). In HT1080-PKO, over-expression of either WNT3A or WNT4 induced cell-autonomous DVL2 and DVL3 phosphorylation (**Fig. 6A**). DVL activation was independent of paracrine signaling, as Wnt over-expression in HT1080-PKO did not activate LRP6 or induce *AXIN2* expression. Interestingly, this cell-autonomous DVL activation by Wnt over-expression occurred despite the inability of HT1080 to respond to paracrine WNT4 (**Fig. 3**), and the lack of WNT3A secretion in HT1080-PKO (**Fig. 2**). Similarly, Wnt over-expression did not activate LRP6 in MM134 (**Supplemental Fig. 6**) further supporting that secreted WNT4 does not activate paracrine signaling. These data support that WNT4 and WNT3A can mediate intracellular signaling independent of secretion or activation of extracellular signaling (ie. LRP6).

**Figure 6.**
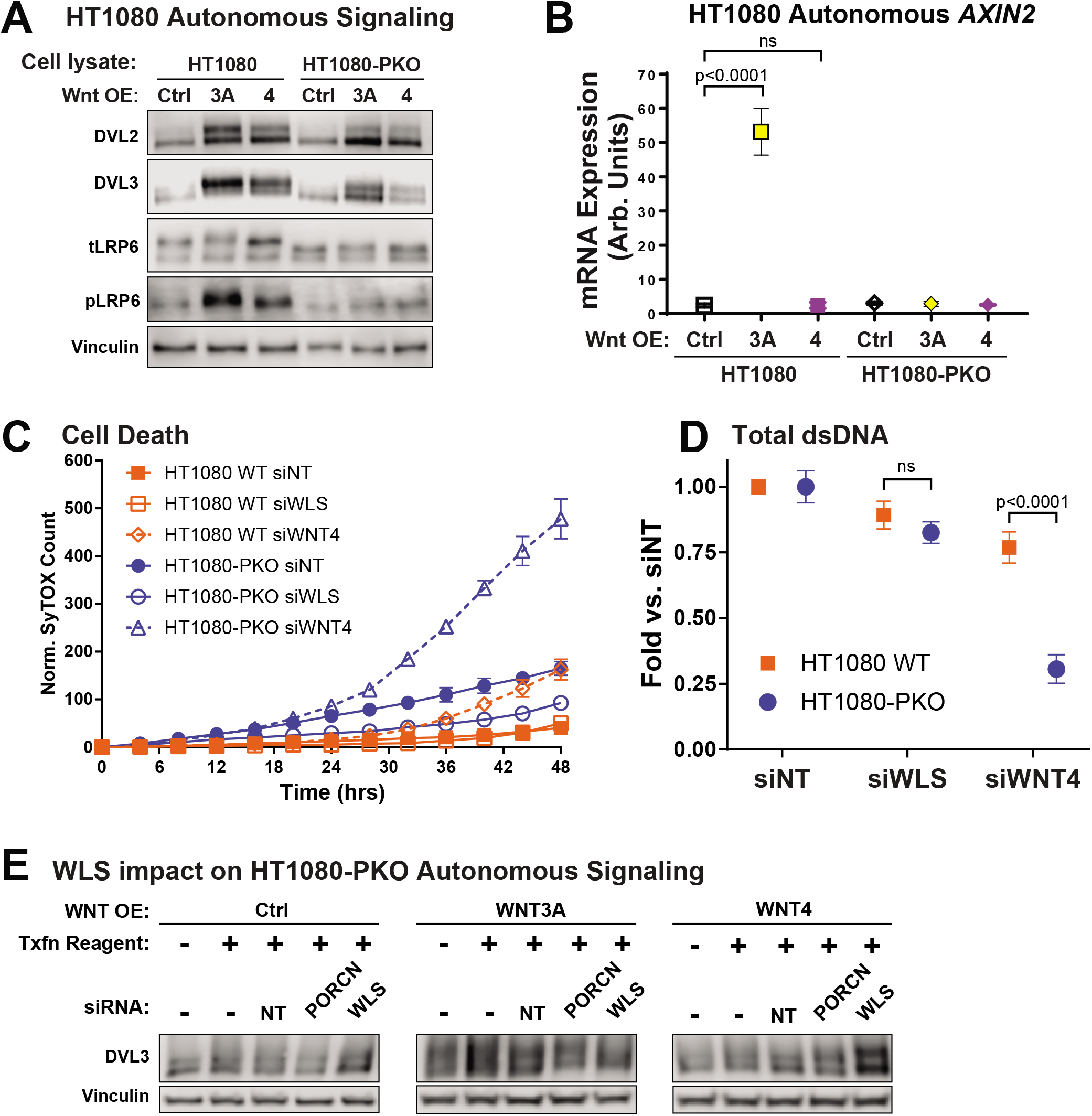
WNT4 and WNT3A have cell autonomous activity independent of PORCN. (A), Whole cell lysates from HT1080 cell lines were harvested and immunoblotted for DVL2, DVL3, total LRP6 and phosphorylated LRP6. (B), mRNA from HT1080 cell lines was extracted and qPCR was performed to determine expression levels of AXIN2. Points represent a mean of 3 technical replicates ±SD. (C-D), HT1080 (WT or PORCN null) cell lines transfected with siControl, siWLS, or siWNT4. (C), Cells were live cell imaged for cell death (SyTOX green) and (D), total double-stranded DNA was measured at timecourse completion. Points represent a mean of 6 biological replicates ±SD. (E), HT1080-PKO cell lines were transfected as indicated. Whole cell lysates were harvested and immunoblotted for DVL3.

As WNT4 is required for bone regeneration and cell proliferation (22), we examined if WNT4 is similarly essential for proliferation and/or viability of HT1080 or HT1080-PKO. We hypothesized that WNT4 might be dispensable in HT1080, due to redundant Wnt signaling. However, without functional PORCN for secretion and paracrine signaling of Wnt family members, HT1080-PKO may become reliant on cell-autonomous PORCN-independent WNT4 signaling. WNT4 knockdown induced ~21% cell death at 48h post-knockdown in HT1080 (**Fig. 6C**), leading to a modest suppression of proliferation (**Fig. 6D**). In contrast, WNT4 knockdown in HT1080-PKO strongly suppressed growth, and cell death was accelerated (~70% cell death at 48h post-knockdown). Importantly, minimal cell death was induced by WLS knockdown in either HT1080 or HT1080-PKO, despite global suppression of Wnt secretion in the latter with ablation of both PORCN and WLS. The sensitivity of HT1080-PKO to knockdown of WNT4, but not WLS, supports a critical role for secretion-independent functions of WNT4, in particular in the absence of paracrine signaling by other Wnt proteins.

To confirm that cell-autonomous PORCN-independent WNT signaling was similarly independent of WLS, we knocked-down WLS or PORCN in HT1080-PKO Wnt over-expressing cells and assessed DVL3 activation (**Fig. 6E**). siPORCN did not affect DVL3 activation, consistent with our above observations. DVL3 activation in HT1080-PKO:W3 was suppressed, suggesting disparate roles of PORCN and WLS in activating paracrine versus cell-autonomous signaling. However, siWLS did not impact cell-autonomous DVL3 activation by WNT4, further supporting a secretion-independent cell-autonomous signaling mechanism for WNT4.

## Discussion

The Wnt modifying enzyme PORCN is commonly described as a gatekeeper for the secretion of Wnt proteins, and thus PORCN inhibition is an approach to broadly block Wnt signaling without targeting cell type-or tissue-specific downstream Wnt pathways. WNT4 signaling is required for survival and proliferation of ILC cells, but we show that PORCN is dispensable, calling into question the role of PORCN in WNT4 signaling. PORCN was not required for WNT4 secretion from a panel of cell lines, as genetic or chemical PORCN blockade had no effect on WNT4 section. However, WNT4 was not capable of activating paracrine Wnt signaling in any model tested, despite the ability of recombinant human WNT4 to do so in a context-dependent manner. These data together suggest that secreted WNT4 may not be responsible for driving signaling in WNT4-expressing cells. Instead, we determined that WNT4 and WNT3A can activate cell-autonomous, intracellular signaling independent of secretion (**Fig. 7, Supplemental Fig. 7**). This unique mode of Wnt signaling is likely essential for the survival and proliferation of WNT4-dependent cells.

**Figure 7.**
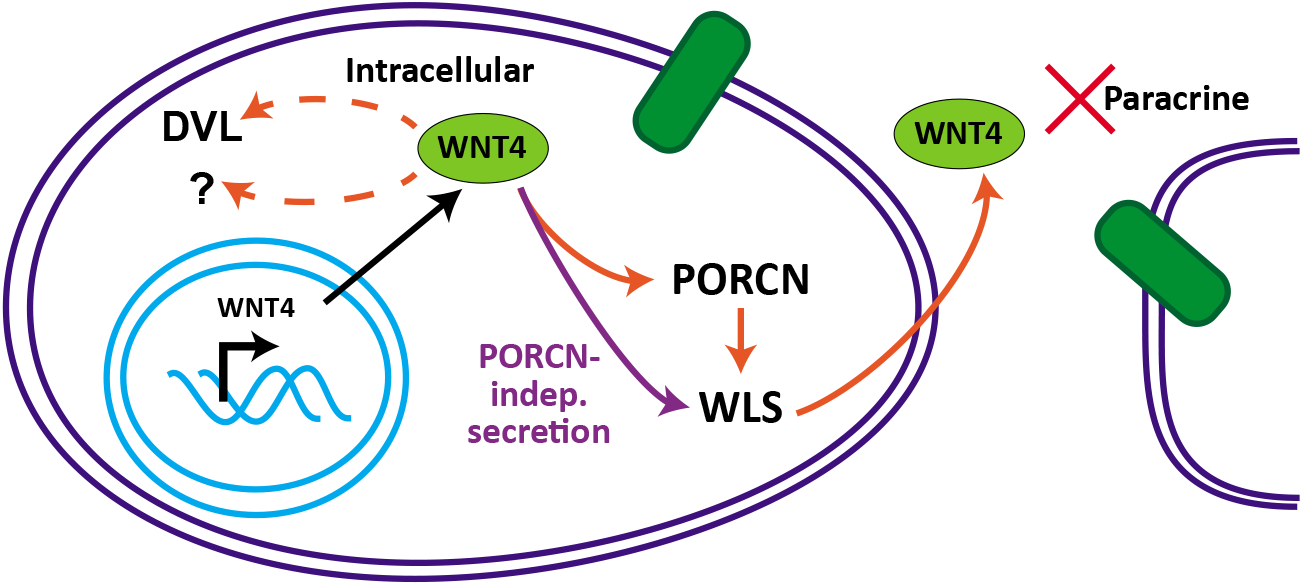
WNT4 activates signaling in an atypical manner independent of secretion. WNT4 can be modified by PORCN during the process of secretion, but WNT4 can instead be secreted in a PORCN-independent, WLS-dependent manner. In either case, secreted WNT4 does not have paracrine activity in any context tested herein. Cell-autonomous WNT4 signaling can activate DVL proteins intracellularly, and likely signals via other pathways in ILC cells.

Our observations of PORCN-independent WNT4 signaling are supported by other studies showing a disconnect between PORCN activity and Wnt secretion/signaling, and support that the requirement for PORCN is context-dependent and not absolute. Nusse and colleagues reported PORCN-independent secretion and activity of *Drosophila* WntD (49). WntD is secreted at high levels in fly tissues and cell culture models independent of both Porcupine and Wntless, and ablation of either Porcupine or Wntless did not affect WntD signaling. WntD utilizes the early secretory pathway protein Rab1 GTPase (RAB1A homolog) for secretion, a distinct secretion mechanism versus Porcupine-mediated secretion of Wingless (WNT1 homolog). WntD lacks the conserved serine residue that is palmitoylated by PORCN (49), unique among Wnt proteins, but this supports that Wnt proteins can be secreted and signal independent of PORCN-driven modification. PORCN-independent Wnt secretion and signaling was also observed in human primary cells by Richards et al (50). Neither PORCNi (IWP-2) nor *PORCN* siRNA knockdown suppressed secretion of any endogenously expressed Wnt proteins from CD8+ T-cells (Wnts 1, 3, 5B, 10B) or astrocytes (Wnts 1, 3, 6, 7A, 10A, 16). Wnt proteins secreted from PORCNi-treated cells were functional in conditioned medium experiments, but PORCN-independent secretory mechanisms were not characterized. Other studies have shown that PORCN may differentially regulate the activity of individual Wnt proteins. In HEK293T cells, over-expression of PORCN and WLS together enhanced secretion of WNT1 but suppressed WNT1-induced paracrine or autocrine activation of β-catenin (versus WLS overexpression alone) (51). This was not observed with WNT3A, indicating PORCN specifically suppressed paracrine activity of WNT1, despite driving WNT1 secretion. Wnt-specific Porcupine functions have also been reported in Zebrafish (52). Knockdown of *porcn* in Zebrafish embryos suppressed secretion of Wnt5a and resulted in defects in non-canonical (β-catenin-independent) Wnt signaling. Conversely, canonical β-catenin-dependent Wnt signaling was not altered by *porcn* knockdown, and secretion of Wnt3a was not impaired. Similar data regarding specifically WNT4 are limited. A clinical *WNT4* mutation (L12P (24), discussed below) blocks WNT4 palmitoylation but not secretion, yet ultimately is associated with WNT4 loss-of-function, consistent with a disconnect between PORCN, WNT4 secretion, and WNT4 signaling. These studies together show Wnt proteins secreted in PORCN-independent manners can be active in paracrine signaling models, but to our knowledge, this is the first report of PORCN-independent Wnt activity that is also independent of secretion. Of note, Kurita et al recently demonstrated that *Wnt4* siRNA in mouse pancreatic β-cells suppressed glucose-induced insulin secretion, but recombinant Wnt4 had no effect on insulin secretion (53). These data parallel our findings and support that WNT4 signals independent of PORCN, secretion, and paracrine activity.

Our data, together with the above reports on PORCN-independent Wnt signaling, highlight that the roles of PORCN and WLS in Wnt modification, secretion, and signaling are Wnt-specific, as well as cell type- and context-dependent. This observed context-dependence includes not only the specific “receiver” cells in question (perhaps best understood in the context of differentially expressed FZD receptors), but also includes the source of the Wnt protein (**Fig. 7, Supplemental Fig. 7**). For example, WNT3A secreted from HT1080 robustly activated Wnt signaling in 3T3-E1 or HT1080-PKO cells, and PORCN was required for both secretion and paracrine activity. In contrast, WNT3A secreted from MM134 cells activated Wnt signaling in 3T3-E1 but not HT1080-PKO cells, and PORCN was required for paracrine activity but not secretion. Cell-type specific Wnt protein post-translational modification (e.g. glycosylation patterns) may drive these differences (51, 54). This parallels our observation that recombinant Wnt proteins were more promiscuously able to activate signaling than secreted Wnt proteins (eg. rWNT4, but not secreted WNT4, activated 3T3-E1 Wnt signaling). rWnt proteins may thus represent a specific functional context versus secreted Wnt protein.

WNT4 and WNT3A likely signal via at least three distinct mechanisms: 1) as a secreted protein with PORCN modification; 2) as a secreted protein without PORCN modification; 3) by a cell-autonomous mechanism independent of secretion. This may explain the myriad context-dependent signaling controlled by WNT4, including activating canonical β-catenin activity (15, 36, 55), repressing β-catenin-driven transcription (56, 57), or activating non-canonical Wnt signaling pathways (22, 58). The signaling differences we observed between recombinant and secreted WNT4 indicate differential protein processing may guide Wnt proteins to activate distinct signaling pathways. The commercial recombinant Wnt proteins used in our study lack the N-terminal signal peptide (residues 1-22 for WNT4), however, Wnt signal peptides may have important roles in the regulation of signaling activity (45). The WNT4 signal peptide has uniquely high content of arginine (14%) and serine (18%) compared to other human Wnts (average 5% and 9%, respectively), whereas WNT3A has no charged or polar residues in its signal peptide. Mutation in the signal peptide (L12P) of WNT4 has also been linked to Mayer-Rokitansky-Küster-Hauser syndrome (24). The L12P mutant functions as a dominant negative inhibitor and suppresses the activity of wild-type WNT4 when co-expressed. Although the L12P mutant protein is not palmitoylated, it is secreted and does not prevent secretion of wild-type protein (24). These observations are consistent with our findings of PORCN-independent WNT4 secretion. The mechanism of dominant-negative activity has not been described but suggests distinct forms of WNT4 may drive cell autonomous Wnt signaling. Characterizing distinct WNT4 species is an important future direction.

ILC is a unique context for paracrine Wnt signaling, as we observed that both WNT3A and WNT4 were secreted from ILC cells in a PORCN-independent manner. While WNT3A activity remained dependent on PORCN (similar to Wnt5a in 293T cells (59)), the role of WNT4 processing in paracrine activity is unclear. However, this atypical Wnt processing in ILC cells (also observed in relation to glycosylation) may be related to broader Wnt signaling dysfunction. The genetic hallmark of ILC is the loss of E-cadherin *(CDH1)* (42), which leads to dysfunction of catenin proteins, including activation of p120 catenin (60) and inactivation of β-catenin. E-cadherin loss in ILC leads to a loss of β-catenin protein in both patient tumors and cell lines (41, 42), and as a result, a β-catenin-driven TOP-Flash reporter cannot be activated in ILC cells (41). Catenin dysfunction was previously postulated as being linked to PORCNi sensitivity, and ILC patients were specifically included in a trial of WNT974 (NCT01351103). This trial opened in 2011, but by 2015 ILC patients were removed from the inclusion criteria. It is unclear whether this is due to accrual problems or a lack of efficacy, as neither have been reported for ILC patients on this trial, although our data suggest PORCNi are unlikely to have clinical efficacy for ILC. This highlights the importance of defining the unique context for Wnt signaling in ILC, in particular for WNT4, based on our prior findings (41). Our laboratory has begun to characterize WNT4-driven signaling in ILC cells, which are likely mediated by PORCN-independent, cell-autonomous WNT4 signaling.

Our study uses diverse cell line models to investigate Wnt secretion and paracrine activity of Wnt proteins. We used non-tagged Wnt expression constructs, eliminating previously described complications caused by epitope tags on Wnt proteins. Wnt signaling activity was measured by complementary pathway readouts that facilitated our study of context-dependent Wnt signaling activities. The cell line models used for Wnt secretion were all cancer-derived, and thus further study is needed to translate our findings to normal tissue, developmental, or *in vivo* contexts. We observed identical processing and secretion for endogenous WNT4 as for over-expressed WNT4 (models herein express low or no endogenous WNT3A), but over-expression was required to facilitate signaling experiments. Future studies will determine the contribution of endogenous Wnt protein levels to activating the signaling pathways discussed herein.

The secretion and paracrine activity of Wnt proteins are heavily context-dependent, as WNT3A and WNT4 present with differing dependence on PORCN for secretion and paracrine activity in distinct model systems. Secretion of WNT4 is PORCN-independent, but WNT4 did not present paracrine activity in cells that were dependent on WNT4, indicative of secretion-independent, cell-autonomous activity. Both WNT4 and WNT3A presented cell-autonomous activity independent of secretion. Our studies identify a PORCN-independent mode of Wnt signaling may be critical to understanding WNT4-driven cellular contexts or those which are otherwise considered to have dysfunctional Wnt signaling.

## Methods and Materials

### Cell culture

HT1080 and PORCN-knockout HT1080 (HT1080-PKO; clone delta-19) were a generous gift from Dr. David Virshup (43), and were maintained in DMEM/F12 (Corning, Corning, NY, USA, cat#10092CV) + 10% FBS (Nucleus Biologics, San Diego, CA, USA, cat#FBS1824). MDA MB 134VI (MM134; ATCC, Manassas, VA, USA) and SUM44PE (BioIVT, Westbury, NY, USA) were maintained as described (40).

PEO1 and HCC1428 (ATCC) were maintained in DMEM/F12 + 10% FBS. MC3T3-E1 (ATCC) were maintained in MEM Alpha without ascorbic acid (Thermo Fisher Scientific, Waltham, MA, USA, cat#A10490-01) + 10% FBS. Wnt overexpression lines were generated by lentiviral transduction of Wnt expression plasmids (see below) with selection of antibiotic-resistant pools, and were maintained in 2.5 μg/mL blasticidin. PEO1:W3 were established by us previously (61). All lines were incubated at 37°C in 5% CO_2_. Cell lines are authenticated annually by the University of Arizona Genetics Core cell line authentication service and confirmed to be mycoplasma negative every four months. Authenticated cells were in continuous culture <6 months.

### Reagents and plasmids

LGK974 was obtained from Cayman Chemical (Ann Arbor, MI, USA, cat#14072) and was dissolved in DMSO. WntC59 was obtained from Tocris Biosciences (Bristol, UK, cat#5148) and was dissolved in DMSO. Fulvestrant was obtained from Tocris Biosciences (#1047) and was dissolved in EtOH. Tunicamycin was obtained from Cayman Chemical (# 11445) and reconstituted in DMSO. Recombinant human WNT3A (#5036-WN-010) and WNT4 (#6076-WN-005) were obtained from R&D Systems (Minneapolis, MN, USA) and reconstituted per the manufacturer’s instructions.

Wnt plasmids used in this publication were a gift from Drs. Marian Waterman, David Virshup and Xi He (Addgene, Watertown, MA, USA, kit #1000000022) (8). The M50 Super 8x TOPFlash plasmid was a generous gift from Dr. Randall Moon (Addgene, plasmid #12456) ((62). Lentiviral vectors for WNT3A and WNT4 were generated by Gateway Recombination of pENTR-STOP Wnt open reading frames to pLX304, a kind gift from Dr. Bob Sclafani.

### Transient transfection assays

HT1080 cells were transfected with Active WNT3A-V5 (G-8) and Active WNT4-V5 (G-10) from Addgene kit #1000000022. The transfection used Lipofectamine LTX Reagent with PLUS Reagent (Thermo Fisher Scientific, Cat# 15338100) using the manufacturer’s instructions.

### Protein extraction from conditioned medium

HT1080, MM134, HCC1428, or PEO1cells were plated in full medium (as above, 10% FBS). Decreased FBS is required to reduce competition of serum proteins with secreted Wnt proteins during extraction. 24hrs post-plating, medium was changed to reduced serum medium: DMEM/F12 + 2% FBS (HT1080, PEO1); DMEM + 2% FBS (HCC1428); DMEM/L15 + 5% FBS (MM134). Conditioned medium was harvested 4-5 days later for HT1080 cells and 6-7 days later for MM134, HCC1428 and PEO1 cells, typically once medium acidification was apparent by phenol red color change. SUM44PE cells are normally cultured in low serum (DMEM/F12 + 2% CSS), so standard medium was allowed to condition for 6-7 days before harvesting. Medium was centrifuged at 300xg for 4min to pellet any cells or debris. The supernatant was then syringe-filtered using a 0.2μm filter. The protein concentration of the conditioned medium was measured using a Pierce BCA Protein Assay Kit (Thermo Fisher Scientific, cat#23225), and medium volumes, normalized to total protein, were adjusted with sterile-filtered PBS. Strataclean resin (Agilent Technologies, Santa Clara, CA, USA, cat#400714) was added to the conditioned media at a ratio of 10μL of resin to 100μg of medium protein, and vortexed to re-suspend the resin. The medium+resin mixture was incubated, rotating at 4C for 30min, then centrifuged at 425xg for 1min at 4C. The supernatant was then aspirated and the resin was washed using sterile-filtered PBS. The resin was centrifuged again at 425xg for 1min at 4C, and the supernatant was aspirated. An equal volume of 2X Laemmli sample buffer (Bio-Rad Laboratories, Hercules, CA, USA, cat#1610737) was added to the resin to release bound protein, and the slurry was heated at 95C for 5min. Slurry equivalent to 100ug of conditioned medium protein was run on SDS-PAGE gels to detect secreted WNT3A or WNT4 via immunoblotting (below).

### Alkaline phosphatase assay

MC3T3-E1 cells were seeded into a 96-well plate at 10,000 cells/well. 24hrs later the cells were treated with either conditioned media (CM) or recombinant WNTs (rWNTs). 72hrs later, alkaline phosphatase activity was assessed, using *para*-Nitrophenyl phosphate (pNp) as a substrate and measuring absorbance at 405nm. Assay protocol is based on Nakamura et al (63) with buffer modifications provided by the recombinant Wnt manufacturer (R&D Systems). A more detailed protocol is available upon request. para-Nitrophenyl Phosphate tablets were obtained from Cayman Chemical (cat# 400090).

### Co-culture and Dual Luciferase Assay

HT1080-PKO cells were transfected with both a TOP-FLASH and renilla reporter plasmid. 24 hours later, these cells were co-cultured with either HT1080 or HT1080-PKO WNT overexpressing cells, with or without 10nM LGK974 treatment. 48 hours later, luciferase and renilla activity was assessed using Promega’s Dual Luciferase Assay (Promega, Madison, WI, USA, cat# E1910) and a BioTek Synergy 2 microplate reader (BioTek, Winooski, VT, USA) with a dual injector system according to the manufacturer’s instructions.

### Proliferation and viability assays

Cells were seeded in a 96-well plate and 24hrs later cells were treated with 100nM Sytox green (Thermo Fisher Scientific, cat# S7020). The plate was then placed into an Incucyte Zoom (Essen Bioscience, Ann Arbor, MI, USA) for 4-5 days where they were imaged every 4hrs at 10x magnification. Cell confluence and cell death (Sytox green-positive counts) were assessed using Incucyte S3 software (v2018A). After time-course completion, total double-stranded DNA was measured by hypotonic lysis of cells in ultra-pure H_2_O, followed by addition of Hoechst 33258 (Thermo Fisher Scientific, #62249) at 1μg/mL in Tris-NaCl buffer (10mM Tris, 2M NaCl; pH 7.4) at equivalent volume to lysate. Fluorescence (360nm ex / 460nm em) was measured on a Bio-Tek Synergy 2 microplate reader.

### RNA interference

siRNAs were reverse transfected using RNAiMAX (ThermoFisher) according to the manufacturer’s instructions. All constructs are siGENOME SMARTpool siRNAs (GE Healthcare Dharmacon, Lafayette, CO, USA): Non-targeting pool #2 (D-001206-14-05), Human *WNT4* (M-008659-03-0005), Human *PORCN* (M-009613-00) and Human *WLS* (M-018728-01-0005). Details regarding validation of the specific effects of the *WNT4* siRNA pool are previously described (41).

### Gene expression analyses

RNA extractions were performed using the RNeasy Mini kit (Qiagen, Venlo, Netherlands); mRNA was converted to cDNA on an Eppendorf Mastercycler Pro (Eppendorf, Hamburg, Germany) and using Promega reagents: Oligo (dT)_15_ primer (cat# C110A), Random Primers (cat# C118A), GoScript 5x Reaction Buffer (cat# A500D), 25mM MgCl2 (cat# A351H), 10mM dNTPs (cat# U1511), RNasin Plus RNase Inhibitor (cat# N261B) and GoScript Reverse Transcriptase (cat# A501D). qPCR reactions were performed with PowerUp SYBR Green Master Mix (Life Technologies, cat # 100029284) on a QuantStudio 6 Flex Real-Time PCR system. Expression data were normalized to *RPLP0.* The following primers were used: *RPLP0,* Forward – CAGCATCTACAACCCTGAAG, Reverse – GACAGACACTGGCAACATT; *WNT4,* Forward – GCCATTGAGGAGTGCCAGTA, Reverse – CCACACCTGCCGAAGAGATG; *WNT3A,* Forward – ATGGTGTCTCGGGAGTT, Reverse – TGGCACTTGCACTTGAG; *PORCN,* Forward – ACCATCCTCATCTACCTACTC, Reverse – CCTTCATGGCCACAATCA; *AXIN2:* Forward – CTCTGGAGCTGTTTCTTACTG, Reverse – CTCTGGAGCTGTTTCTTACTG.

### Immunoblotting

Whole-cell lysates were obtained by incubating cells in lysis buffer (1% Triton X-100, 50mM HEPES pH 7.4, 140mM NaCl, 1.5mM MgCl_2_, 1mM EGTA, 10% glycerol; supplemented with Roche protease/phosphatase inhibitor cocktail (Sigma-Aldrich, St. Louis, MO, USA)) for 30min on ice. Cells were centrifuged at 16000xg for 15min at 4C and the resulting supernatant was collected for analysis. Protein concentration was measured using the Pierce BCA Protein Assay Kit (#23225). Standard methods were used to perform SDS-PAGE. Proteins were transferred onto PVDF membranes. Antibodies were used according to manufacturer’s recommendations: WNT4 (R&D, MAB4751, clone# 55025, 1:1,000), WNT3A (R&D, MAB13242, clone# 217804.2R, 1:1,000), DVL2 (Cell Signaling Technology, Danvers, MA, USA, cat#3216, 1:2,000), DVL3 (Cell Signaling, 3218, 1:2,000), pLRP6 (s1490, Cell Signaling, 2568, 1:2,000), LRP6 (Cell Signaling, 2560, clone# C5C7, 1:2,000), Vinculin (Cell Signaling, cat# 13901S 1:10,000) and V5 (Novus Biologicals, Centennial, CO, USA, NB100-62264, clone #SV5-PK1, 1:2,500). Specificity of WNT4 antibody was validated using siWNT4 (**Figure S3**) and the recognition of recombinant WNT4 compared to endogenous and overexpressed WNT4 (**Figure 4E-F**). Secondary antibodies were used according to manufacturer’s instruction and were obtained from Jackson ImmunoResearch Laboratories (West Grove, PA, USA): Goat Anti-Mouse IgG (cat # 115-035-068), Goat Anti-Rabbit IgG (cat# 111-035-045) and Goat Anti-Rat IgG (cat# 112-035-062). All secondary antibodies were used at a dilution of 1:10,000. Chemiluminescence was used to detect antibodies and either film or the LI-COR c-Digit (LI-COR Biosciences, Lincoln, NE, USA) was used to develop the immunoblots. Of note, WNT4 MAB4751 detects a prominent non-specific band at ~50kD in immunoblots from cell lysates. This non-specific target largely precludes detection of WNT4, but cutting immunoblot membranes immediately below a 50kD ladder marker prevents this issue. This non-specific band was not detected in WNT4 immunoblots from conditioned medium. Similarly, in immunoblots of conditioned medium WNT3A MAB13242 detects a prominent non-specific band at ~60kD that precludes detection of secreted WNT3A; cutting membranes above a 50kD ladder marker prevents this issue. This non-specific band was not detected in WNT3A immunoblots from cell lysates.

### Statistical Considerations

Prism was used for all graphical representation and statistical analyses. All western blot figures are representative of at least two to three independent experiments.

### Software

Prism (GraphPad Software, San Diego, CA, USA, version 7.02) and Image Studio (LI-COR Biosciences, Lincoln, NE, USA, version 5.2) were used to obtain and analyze that data presented in this manuscript.

## Acknowledgements

The authors thank Dr. Caroline Alexander (U. Wisconsin) for critical reading of the manuscript.

## Data availability

Data associated with experiments herein will be available at an Open Science Framework repository (64) (https://doi.org/10.17605/OSF.IO/7X8NG).

## List of Abbreviations

AP: Alkaline Phosphatase
AXIN2: Axin related protein 2
CDH1: E-cadherin
CM: Conditioned Media
DVL: Disheveled
Fulv: Fulvestrant
FZD: Frizzled
ILC: Invasive Lobular Carcinoma
LRP: Lipoprotein receptor-related protein
PKO: Porcupine knockout
PORCN: O-acyltransferase Porcupine
PORCNi: Porcupine inhibitor
rWNT: Recombinant WNT
sWNT: Secreted WNT
WLS: Wntless
WNT: Wingless/Integrated

---

**Supplemental Figure 1.**
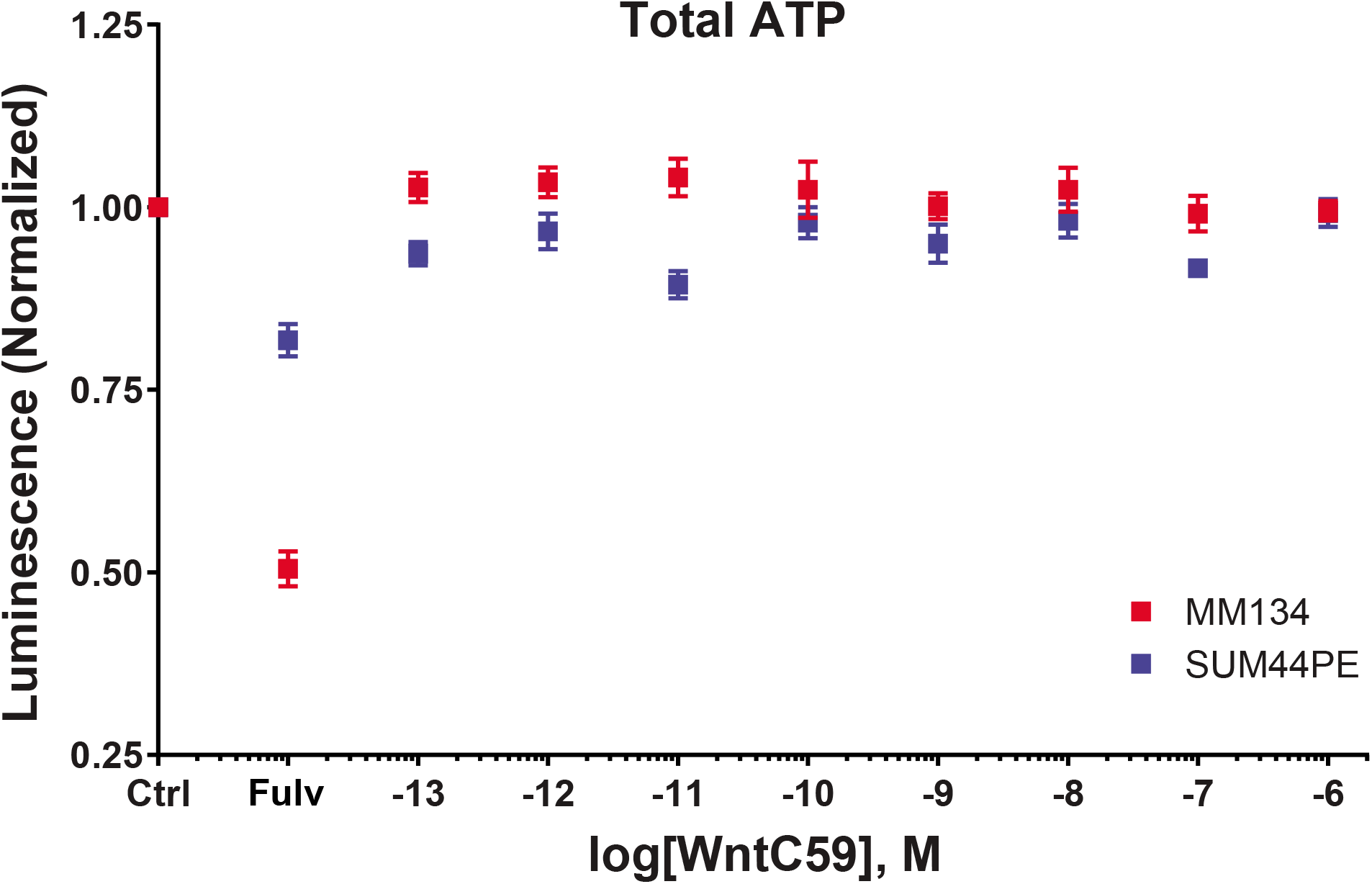
PORCNi (WntC59) does not impact ILC growth. MM134 and SUM44PE were plated and 24hrs later started treatment with either anti-estrogen fulvestrant (Fulv) or increasing concentrations of WntC59. At the timecourse completion of either 5 or 7 days, total ATP was measured using Promega Cell-Titer Glo. Points represent the mean of 6 biological replicates + SD.

**Supplemental Figure 2.**
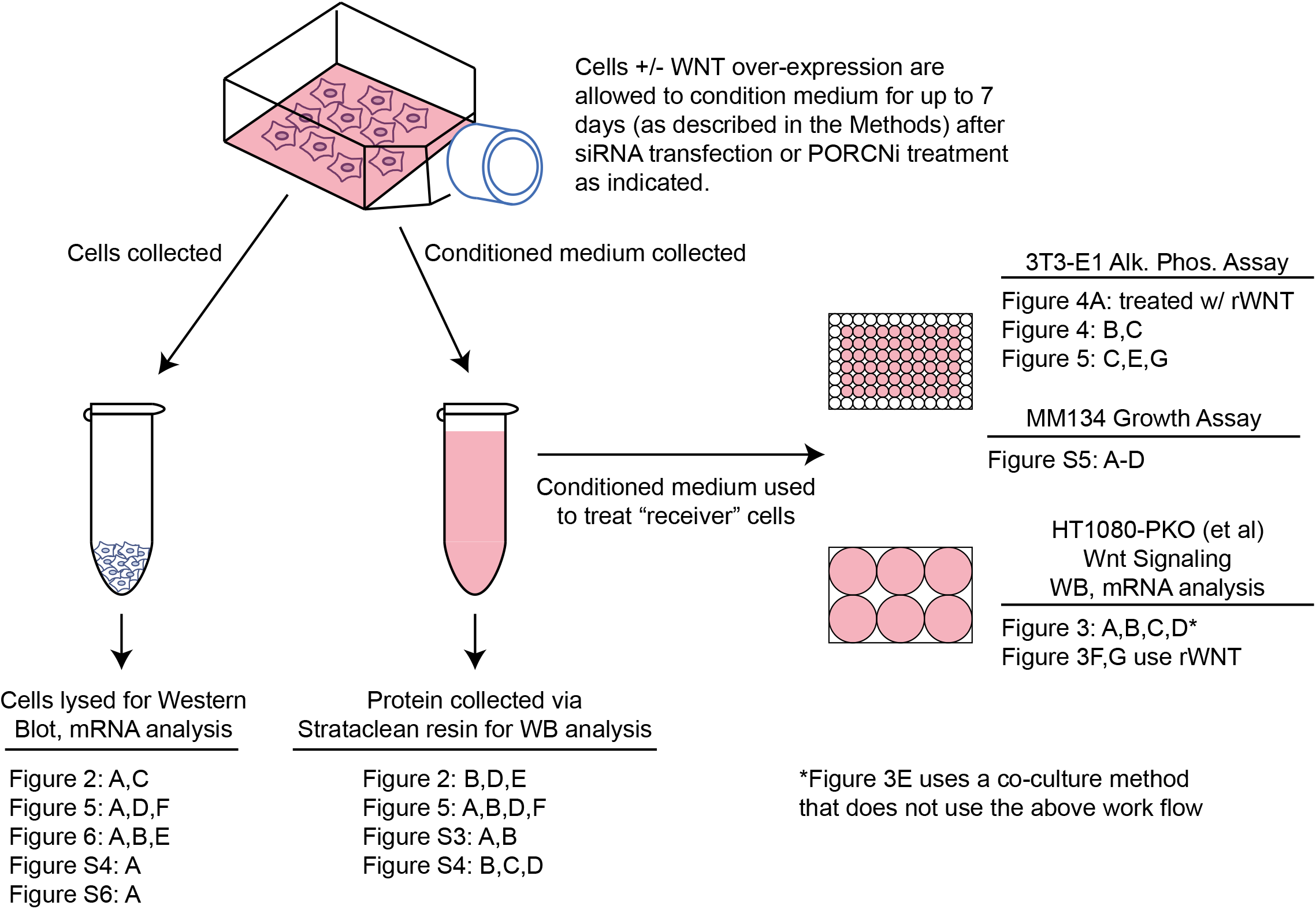
Experimental Workflow for Assessing Wnt Secretion and Function. Diagram of the workflow for experiments aimed at assessing the ability of Wnt proteins to be secreted and their functionality. Experimental endpoints are labeled with corresponding figures.

**Supplemental Figure 3.**
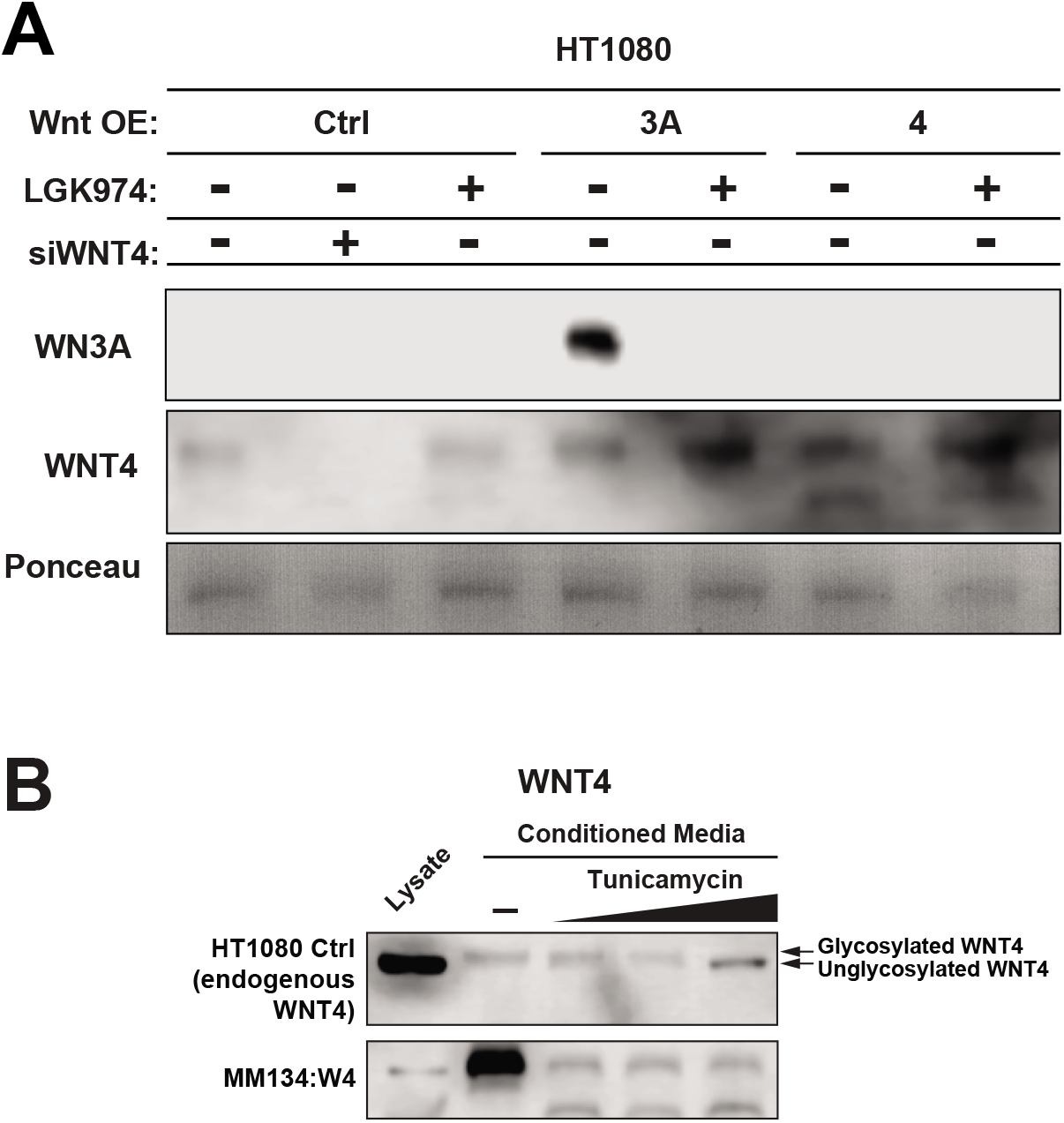
WNT4 secretion is PORCN-independent, but secreted WNT4 is post-translationally modified. **(A),** HT1080 cells were reverse transfected with siWNT4. 24hrs later media was changed and cells were treated with or without LGK974 (10nM). Conditioned media (CM) was harvested 5 days later and total protein was collected from immunoblotting (as described in the Methods and Materials) for WNT3A or WNT4. **(B),** Conditioned media was harvested from either HT1080:WT or MM134:Wnt4 over-expressing cells after treatment (3 or 5 days respectively) with increasing concentrations of tunicamycin (0.5uM, 1uM or 10uM). WNT4 presence was detected from untreated whole cell lysate and conditioned media.

**Supplemental Figure 4.**
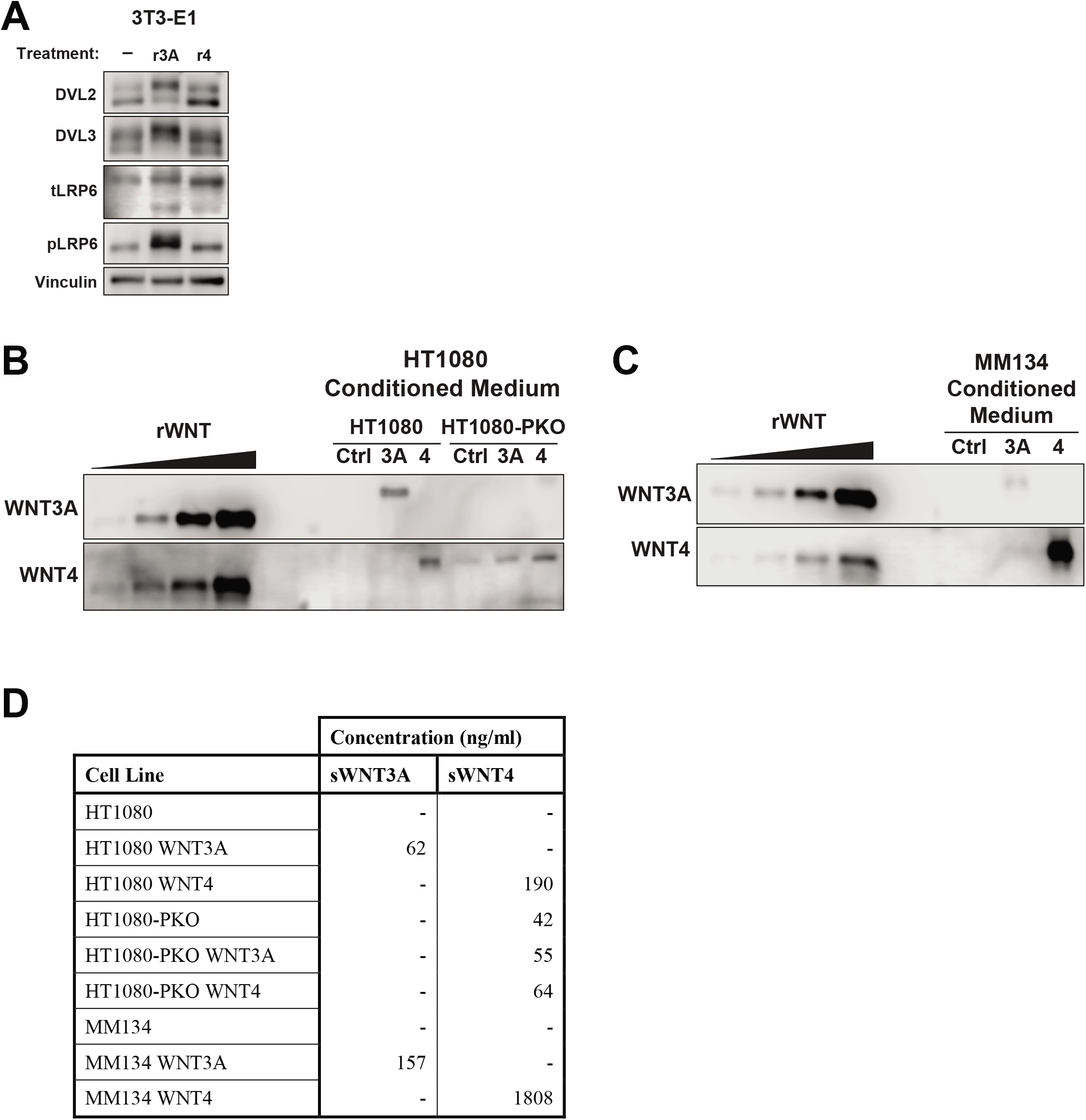
Secreted Wnt proteins are at concentrations sufficient to activate paracrine signaling. (A), 3T3-E1 cells were treated for 24hrs with 300ng/ml of either rWNT3A or rWNT4. Immunoblots of whole cell lysates were probed for DVL2, DVL3, total LRP6 and phosphorylated LRP6. (E-F), Serial dilutions of rWNTs (2.5, 5, 10, 20ng) were run on an immunoblot alongside CM collected from HT1080 or MM134 cell lines. Blots were probed for either WNT3A or WNT4, respectively. (G), Estimated concentrations of secreted WNTs (sWNT) present in CM based on linear regression of rWNTs. Volume of CM loaded per well varied from 25-70uL, normalized based on total protein concentration (see Materials and Methods).

**Supplemental Figure 5.**
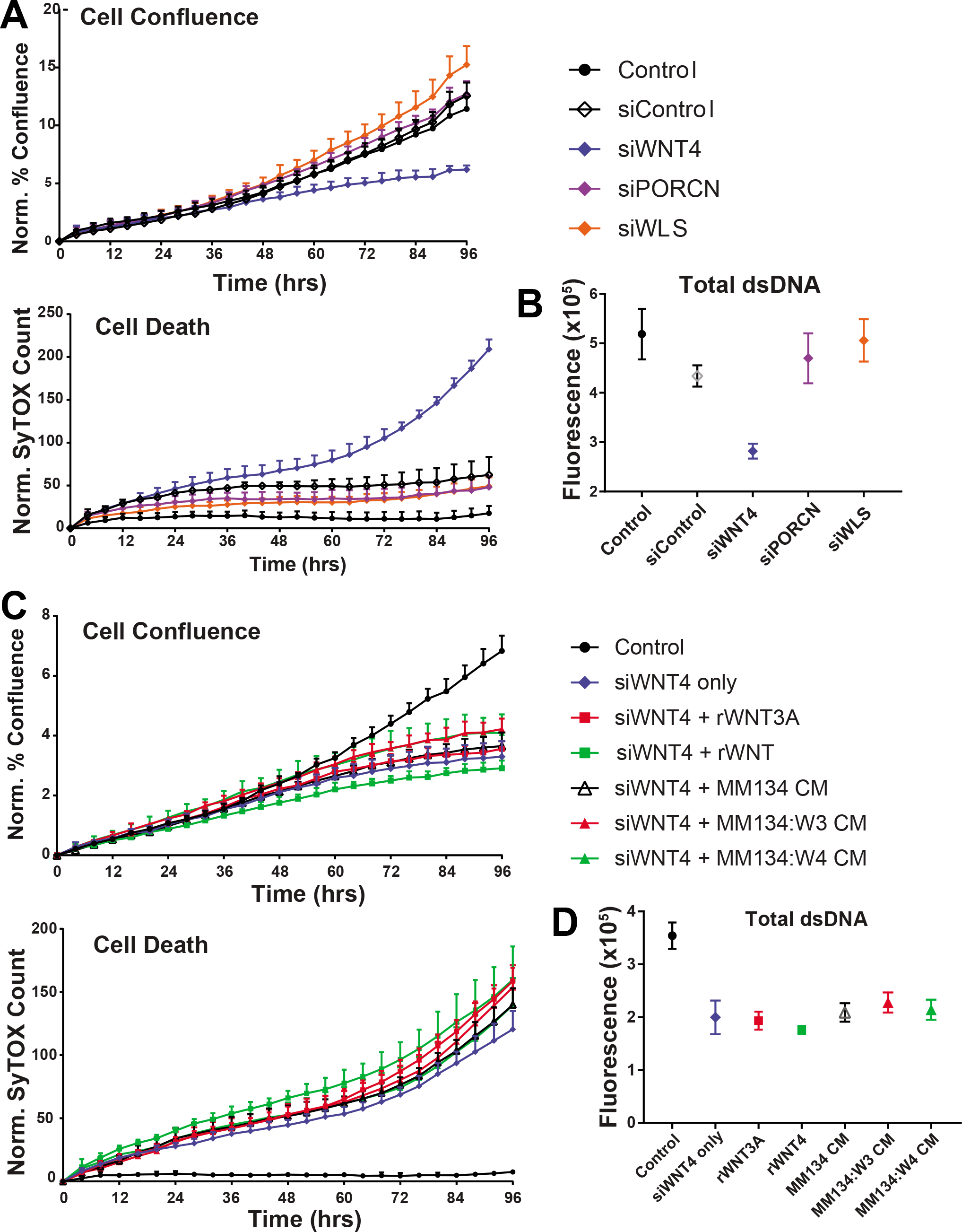
siWLS does not phenocopy siWNT4 and CM does not rescue siWNT4 in MM134. **(A-B),** MM134 were reverse transfected with siRNA as indicated. **(C-D),** 24hrs after WNT4 knockdown, cells were treated with either rWNT3A (62.5mg/ml), rWNT4 (250ng/ml) or conditioned media (CM). The cells were live cell imaged for **(A,C),** proliferation (phase-contrast confluence) and death (SyTOX green incorporation). **(B,D),** Total double stranded DNA was neasured from assays in **(A,C)** at timecourse completion. Point represent mean of six biological replicates ± SD.

**Supplemental Figure 6.**
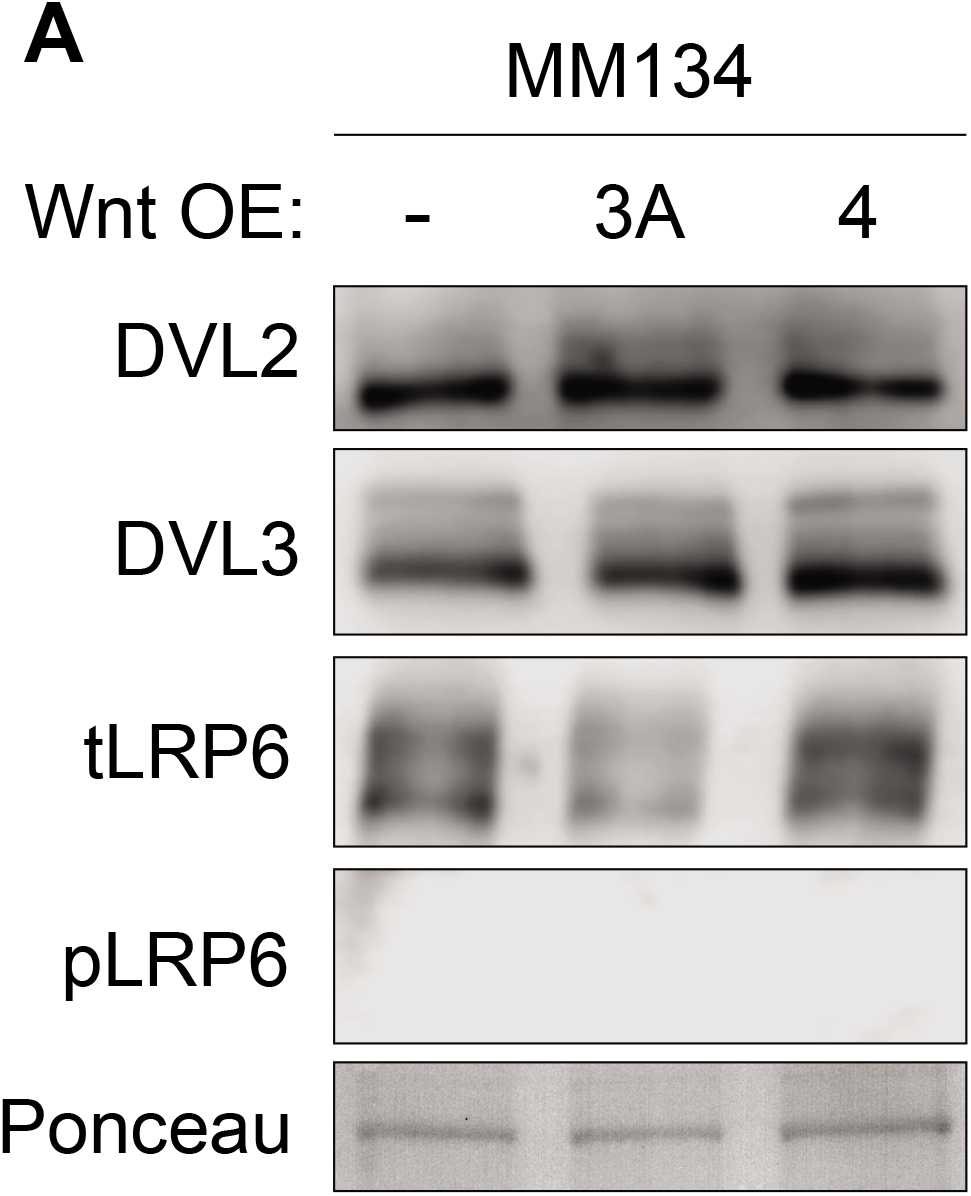
WNT overexpression does not induce autonomous phosphorylation of DVL2/3 in MM134. Whole cell lysates from MM134 were harvested and immunoblotted for DVL2, DVL3, total and phosphor-ylated LRP6.

**Supplemental Figure 7.**
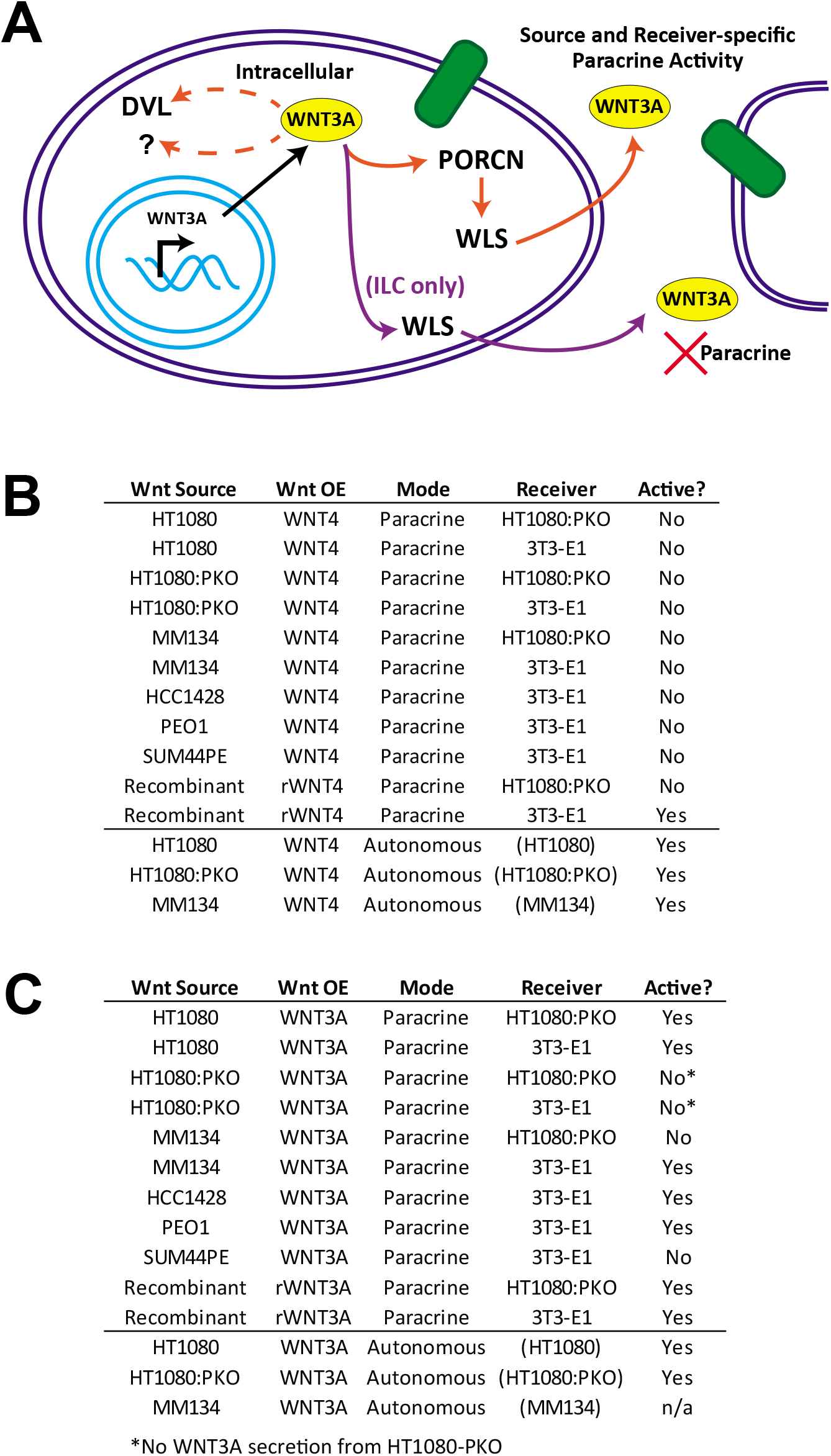
Summary of paracrine Wnt activity per source and receiver context. (A), WNT3A can be secreted independent of PORCN from ILC cells, but PORCN function is overall required for paracrine activity. However, paracrine activity of WNT3A was dependent on the Wnt source and the receiver cell context. (B-C), Summary of results herein for (B) WNT4 and (C) WNT3A.

## References

1. Nusse R, Clevers H. 2017. Wnt/β-Catenin Signaling, Disease, and Emerging Therapeutic Modalities. Cell 169:985–999.

2. Madan B, Ke Z, Harmston N, Ho SY, Frois AO, Alam J, Jeyaraj DA, Pendharkar V, Ghosh K, Virshup IH, Manoharan V, Ong EHQ, Sangthongpitag K, Hill J, Petretto E, Keller TH, Lee MA, Matter A, Virshup DM. 2016. Wnt addiction of genetically defined cancers reversed by PORCN inhibition. Oncogene 35:2197–207.

3. Liu J, Pan S, Hsieh MH, Ng N, Sun F, Wang T, Kasibhatla S, Schuller AG, Li AG, Cheng D, Li J, Tompkins C, Pferdekamper A, Steffy A, Cheng J, Kowal C, Phung V, Guo G, Wang Y, Graham MP, Flynn S, Brenner JC, Li C, Villarroel MC, Schultz PG, Wu X, McNamara P, Sellers WR, Petruzzelli L, Boral AL, Seidel HM, McLaughlin ME, Che J, Carey TE, Vanasse G, Harris JL. 2013. Targeting Wnt-driven cancer through the inhibition of Porcupine by LGK974. Proc Natl Acad Sci U S A 110:20224–9.

4. van Amerongen R. 2012. Alternative Wnt pathways and receptors. Cold Spring Harb Perspect Biol 4:a007914–a007914.

5. Loh KM, van Amerongen R, Nusse R. 2016. Generating Cellular Diversity and Spatial Form: Wnt Signaling and the Evolution of Multicellular Animals. Dev Cell 38:643–55.

6. Brisken C, Hess K, Jeitziner R. 2015. Progesterone and Overlooked Endocrine Pathways in Breast Cancer Pathogenesis. Endocrinology 156:3442–50.

7. Miranda M, Galli LM, Enriquez M, Szabo LA, Gao X, Hannoush RN, Burrus LW. 2014. Identification of the WNT1 residues required for palmitoylation by Porcupine. FEBS Lett 588:4815–24.

8. Najdi R, Proffitt K, Sprowl S, Kaur S, Yu J, Covey TM, Virshup DM, Waterman ML. 2012. A uniform human Wnt expression library reveals a shared secretory pathway and unique signaling activities. Differentiation 84:203–13.

9. Takada R, Satomi Y, Kurata T, Ueno N, Norioka S, Kondoh H, Takao T. 2006. Monounsaturated Fatty Acid Modification of Wnt Protein: Its Role in Wnt Secretion 791–801.

10. Bänziger C, Soldini D, Schütt C, Zipperlen P, Hausmann G, Basler K. 2006. Wntless, a conserved membrane protein dedicated to the secretion of Wnt proteins from signaling cells. Cell 125:509–22.

11. Coombs GS, Yu J, Canning CA, Veltri CA, Covey TM, Cheong JK, Utomo V, Banerjee N, Zhang ZH, Jadulco RC, Concepcion GP, Bugni TS, Harper MK, Mihalek I, Jones CM, Ireland CM, Virshup DM. 2010. WLS-dependent secretion of WNT3A requires Ser209 acylation and vacuolar acidification. J Cell Sci 123:3357–67.

12. Proffitt KD, Madan B, Ke Z, Pendharkar V, Ding L, Lee MA, Hannoush RN, Virshup DM. 2013. Pharmacological inhibition of the Wnt acyltransferase PORCN prevents growth of WNT-driven mammary cancer. Cancer Res 73:502–7.

13. Hayashi M, Baker A, Goldstein SD, Albert CM, Jackson KW, McCarty G, Kahlert UD, Loeb DM. 2017. Inhibition of porcupine prolongs metastasis free survival in a mouse xenograft model of Ewing sarcoma. Oncotarget 8:78265–78276.

14. Agarwal P, Zhang B, Ho Y, Cook A, Li L, Mikhail FM, Wang Y, McLaughlin ME, Bhatia R. 2017. Enhanced targeting of CML stem and progenitor cells by inhibition of porcupine acyltransferase in combination with TKI. Blood 129:1008–1020.

15. Li Q, Kannan A, Das A, Demayo FJ, Hornsby PJ, Young SL, Taylor RN, Bagchi MK, Bagchi IC. 2013. WNT4 acts downstream of BMP2 and functions via β-catenin signaling pathway to regulate human endometrial stromal cell differentiation. Endocrinology 154:446–57.

16. Itäranta P, Chi L, Seppänen T, Niku M, Tuukkanen J, Peltoketo H, Vainio S. 2006. Wnt-4 signaling is involved in the control of smooth muscle cell fate via Bmp-4 in the medullary stroma of the developing kidney. Dev Biol 293:473–83.

17. Tomizuka K, Horikoshi K, Kitada R, Sugawara Y, Iba Y, Kojima A, Yoshitome A, Yamawaki K, Amagai M, Inoue A, Oshima T, Kakitani M. 2008. R-spondin1 plays an essential role in ovarian development through positively regulating Wnt-4 signaling. Hum Mol Genet 17:1278–91.

18. Chang J, Sonoyama W, Wang Z, Jin Q, Zhang C, Krebsbach PH, Giannobile W, Shi S, Wang C-Y. 2007. Noncanonical Wnt-4 signaling enhances bone regeneration of mesenchymal stem cells in craniofacial defects through activation of p38 MAPK. J Biol Chem 282:30938–48.

19. Prunskaite-Hyyryläinen R, Skovorodkin I, Xu Q, Miinalainen I, Shan J, Vainio SJ. 2016. Wnt4 coordinates directional cell migration and extension of the Müllerian duct essential for ontogenesis of the female reproductive tract. Hum Mol Genet 25:1059–73.

20. Strochlic L, Falk J, Goillot E, Sigoillot S, Bourgeois F, Delers P, Rouvière J, Swain A, Castellani V, Schaeffer L, Legay C. 2012. Wnt4 participates in the formation of vertebrate neuromuscular junction. PLoS One 7:e29976.

21. Caprioli A, Villasenor A, Wylie LA, Braitsch C, Marty-Santos L, Barry D, Karner CM, Fu S, Meadows SM, Carroll TJ, Cleaver O. 2015. Wnt4 is essential to normal mammalian lung development. Dev Biol 406:222–34.

22. Yu B, Chang J, Liu Y, Li J, Kevork K, Al-Hezaimi K, Graves DT, Park N-H, Wang C-Y. 2014. Wnt4 signaling prevents skeletal aging and inflammation by inhibiting nuclear factor-κB. Nat Med 20:1009–17.

23. Bernard P, Harley VR. 2007. Wnt4 action in gonadal development and sex determination. Int J Biochem Cell Biol 39:31–43.

24. Philibert P, Biason-Lauber A, Rouzier R, Pienkowski C, Paris F, Konrad D, Schoenle E, Sultan C. 2008. Identification and functional analysis of a new WNT4 gene mutation among 28 adolescent girls with primary amenorrhea and müllerian duct abnormalities: a French collaborative study. J Clin Endocrinol Metab 93:895–900.

25. Vainio S, Heikkilä M, Kispert A, Chin N, McMahon AP. 1999. Female development in mammals is regulated by Wnt-4 signalling. Nature 397:405–9.

26. Biason-Lauber A, De Filippo G, Konrad D, Scarano G, Nazzaro A, Schoenle EJ. 2007. WNT4 deficiency--a clinical phenotype distinct from the classic Mayer-Rokitansky-Kuster-Hauser syndrome: a case report. Hum Reprod 22:224–9.

27. Powell JE, Fung JN, Shakhbazov K, Sapkota Y, Cloonan N, Hemani G, Hillman KM, Kaufmann S, Luong HT, Bowdler L, Painter JN, Holdsworth-Carson SJ, Visscher PM, Dinger ME, Healey M, Nyholt DR, French JD, Edwards SL, Rogers PAW, Montgomery GW. 2016. Endometriosis risk alleles at 1p36.12 act through inverse regulation of CDC42 and LINC00339. Hum Mol Genet 25:5046–5058.

28. Rahmioglu N, Macgregor S, Drong AW, Hedman ÅK, Harris HR, Randall JC, Prokopenko I, Nyholt DR, Morris AP, Montgomery GW, Missmer SA, Lindgren CM, Zondervan KT, Collins FS, International Endogene Consortium (IEC) TGC, Nyholt DR, Morris AP, Montgomery GW, Missmer SA, Lindgren CM, Zondervan KT. 2015. Genome-wide enrichment analysis between endometriosis and obesity-related traits reveals novel susceptibility loci. Hum Mol Genet 24:1185–1199.

29. Mafra F, Catto M, Bianco B, Barbosa CP, Christofolini D. 2015. Association of WNT4 polymorphisms with endometriosis in infertile patients. J Assist Reprod Genet 32:1359–64.

30. Zhang G, Feenstra B, Bacelis J, Liu X, Muglia LJLMLJ, Juodakis J, Miller DE, Litterman N, Jiang P-P, Russell L, Hinds DA, Hu Y, Weirauch MT, Chen X, Chavan AR, Wagner GP, Pavličev M, Nnamani MC, Maziarz J, Karjalainen MK, Rämet M, Sengpiel V, Geller F, Boyd HA, Palotie A, Momany A, Bedell B, Ryckman KK, Huusko JM, Forney CR, Kottyan LC, Hallman M, Teramo K, Nohr EA, Davey Smith G, Melbye M, Jacobsson B, Muglia LJLMLJ. 2017. Genetic Associations with Gestational Duration and Spontaneous Preterm Birth. N Engl J Med 377:1156–1167.

31. Zmuda JM, Yerges-Armstrong LM, Moffett SP, Klei L, Kammerer CM, Roeder K, Cauley JA, Kuipers A, Ensrud KE, Nestlerode CS, Hoffman AR, Lewis CE, Lang TF, Barrett-Connor E, Ferrell RE, Orwoll ES, Osteoporotic Fractures in Men (MrOS) Study Group. 2011. Genetic analysis of vertebral trabecular bone density and cross-sectional area in older men. Osteoporos Int 22:1079–90.

32. Zheng Y, Wang C, Zhang H, Shao C, Gao L-H, Li S-S, Yu W-J, He J-W, Fu W-Z, Hu Y-Q, Li M, Liu Y-J, Zhang Z-L. 2016. Polymorphisms in Wnt signaling pathway genes are associated with peak bone mineral density, lean mass, and fat mass in Chinese male nuclear families. Osteoporos Int 27:1805–1815.

33. Kuchenbaecker KB, Ramus SJ, Tyrer J, Lee A, Shen HC, Beesley J, Lawrenson K, McGuffog L, Healey S, Lee JM, Spindler TJ, Lin YG, Pejovic T, Bean Y, Li Q, Coetzee S, Hazelett D, Miron A, Southey M, Terry MB, Goldgar DE, Buys SS, Janavicius R, Dorfling CM, Van Rensburg EJ, Neuhausen SL, Ding YC, Hansen TVO, Jønson L, Gerdes AM, Ejlertsen B, Barrowdale D, Dennis J, Benitez J, Osorio A, Garcia MJ, Komenaka I, Weitzel JN, Ganschow P, Peterlongo P, Bernard L, Viel A, Bonanni B, Peissel B, Manoukian S, Radice P, Papi L, Ottini L, Fostira F, Konstantopoulou I, Garber J, Frost D, Perkins J, Platte R, Ellis S, Godwin AK, Schmutzler RK, Meindl A, Engel C, Sutter C, Sinilnikova OM, Damiola F, Mazoyer S, Stoppa-Lyonnet D, Claes K, De Leeneer K, Kirk J, Rodriguez GC, Piedmonte M, O’Malley DM, De La Hoya M, Caldes T, Aittomäki K, Nevanlinna H, Margriet JC, Rookus MA, Oosterwijk JC, Tihomirova L, Tung N, Hamann U, Isaccs C, Tischkowitz M, Imyanitov EN, Caligo MA, Campbell IG, Hogervorst FBL, Olah E, Diez O, Blanco I, Brunet J, Lazaro C, Pujana MA, Jakubowska A, Gronwald J, Lubinski J, Sukiennicki G, Barkardottir RB, Plante M, Simard J, Soucy P, Montagna M, Tognazzo S, Teixeira MR, Pankratz VS, Wang X, Lindor N, Szabo CI, Kauff N, Vijai J, Aghajanian CA, Pfeiler G, Berger A, Singer CF, Tea MK, Phelan CM, Greene MH, Mai PL, Rennert G, Mulligan AM, Tchatchou S, Andrulis IL, Glendon G, Toland AE, Jensen UB, Kruse TA, Thomassen M, Bojesen A, Zidan J, Friedman E, Laitman Y, Soller M, Liljegren A, Arver B, Einbeigi Z, Stenmark-Askmalm M, Olopade OI, Nussbaum RL, Rebbeck TR, Nathanson KL, Domchek SM, Lu KH, Karlan BY, Walsh C, Lester J, Hein A, Ekici AB, Beckmann MW, Fasching PA, Lambrechts D, Van Nieuwenhuysen E, Vergote I, Lambrechts S, Dicks E, Doherty JA, Wicklund KG, Rossing MA, Rudolph A, Chang-Claude J, Wang-Gohrke S, Eilber U, Moysich KB, Odunsi K, Sucheston L, Lele S, Wilkens LR, Goodman MT, Thompson PJ, Shvetsov YB, Runnebaum IB, Dürst M, Hillemanns P, Dörk T, Antonenkova N, Bogdanova N, Leminen A, Pelttari LM, Butzow R, Modugno F, Kelley JL, Edwards RP, Ness RB, Du Bois A, Heitz F, Schwaab I, Harter P, Matsuo K, Hosono S, Orsulic S, Jensen A, Kjaer SK, Hogdall E, Hasmad HN, Noor Azmi MA, Teo SH, Woo YL, Fridley BL, Goode EL, Cunningham JM, Vierkant RA, Bruinsma F, Giles GG, Liang D, Hildebrandt MAT, Wu X, Levine DA, Bisogna M, Berchuck A, Iversen ES, Schildkraut JM, Concannon P, Weber RP, Cramer DW, Terry KL, Poole EM, Tworoger SS, Bandera E V., Orlow I, Olson SH, Krakstad C, Salvesen HB, Tangen IL, Bjorge L, Van Altena AM, Aben KKH, Kiemeney LA, Massuger LFAG, Kellar M, Brooks-Wilson A, Kelemen LE, Cook LS, Le ND, Cybulski C, Yang H, Lissowska J, Brinton LA, Wentzensen N, Hogdall C, Lundvall L, Nedergaard L, Baker H, Song H, Eccles D, McNeish I, Paul J, Carty K, Siddiqui N, Glasspool R, Whittemore AS, Rothstein JH, McGuire V, Sieh W, Ji BT, Zheng W, Shu XO, Gao YT, Rosen B, Risch HA, McLaughlin JR, Narod SA, Monteiro AN, Chen A, Lin HY, Permuth-Wey J, Sellers TA, Tsai YY, Chen Z, Ziogas A, Anton-Culver H, Gentry-Maharaj A, Menon U, Harrington P, Lee AW, Wu AH, Pearce CL, Coetzee G, Pike MC, Dansonka-Mieszkowska A, Timorek A, Rzepecka IK, Kupryjanczyk J, Freedman M, Noushmehr H, Easton DF, Offit K, Couch FJ, Gayther S, Pharoah PP, Antoniou AC, Chenevix-Trench G. 2015. Identification of six new susceptibility loci for invasive epithelial ovarian cancer. Nat Genet 47:164–171.

34. Brisken C, Heineman A, Chavarria T, Elenbaas B, Tan J, Dey SK, McMahon JA, McMahon AP, Weinberg RA. 2000. Essential function of Wnt-4 in mammary gland development downstream of progesterone signaling. Genes Dev 14:650–4.

35. Alexander CM, Goel S, Fakhraldeen S a., Kim S. 2012. Wnt signaling in mammary glands: plastic cell fates and combinatorial signaling. Cold Spring Harb Perspect Biol 4.

36. Rajaram RD, Buric D, Caikovski M, Ayyanan A, Rougemont J, Shan J, Vainio SJ, Yalcin-Ozuysal O, Brisken C. 2015. Progesterone and Wnt4 control mammary stem cells via myoepithelial crosstalk. EMBO J 34:641–652.

37. Joshi PA, Jackson HW, Beristain AG, Di Grappa MA, Mote PA, Clarke CL, Stingl J, Waterhouse PD, Khokha R. 2010. Progesterone induces adult mammary stem cell expansion. Nature 465:803–807.

38. Meier-Abt F, Milani E, Roloff T, Brinkhaus H, Duss S, Meyer DS, Klebba I, Balwierz PJ, van Nimwegen E, Bentires-Alj M. 2013. Parity induces differentiation and reduces Wnt/Notch signaling ratio and proliferation potential of basal stem/progenitor cells isolated from mouse mammary epithelium. Breast Cancer Res 15:R36.

39. Kim YC, Clark RJ, Pelegri F, Alexander CM. 2009. Wnt4 is not sufficient to induce lobuloalveolar mammary development. BMC Dev Biol 9:55.

40. Sikora MJ, Cooper KL, Bahreini A, Luthra S, Wang G, Chandran UR, Davidson NE, Dabbs DJ, Welm AL, Oesterreich S. 2014. Invasive lobular carcinoma cell lines are characterized by unique estrogen-mediated gene expression patterns and altered tamoxifen response. Cancer Res 74:1463–74.

41. Sikora MJ, Jacobsen BM, Levine K, Chen J, Davidson NE, Lee A V, Alexander CM, Oesterreich S. 2016. WNT4 mediates estrogen receptor signaling and endocrine resistance in invasive lobular carcinoma cell lines. Breast Cancer Res 18:92.

42. Ciriello G, Gatza ML, Beck AH, Wilkerson MD, Rhie SK, Pastore A, Zhang H, McLellan M, Yau C, Kandoth C, Bowlby R, Shen H, Hayat S, Fieldhouse R, Lester SC, Tse GMK, Factor RE, Collins LC, Allison KH, Chen Y-Y, Jensen K, Johnson NB, Oesterreich S, Mills GB, Cherniack AD, Robertson G, Benz C, Sander C, Laird PW, Hoadley KA, King TA, TCGA Research Network, Perou CM. 2015. Comprehensive Molecular Portraits of Invasive Lobular Breast Cancer. Cell 163:506–519.

43. Proffitt KD, Virshup DM. 2012. Precise regulation of porcupine activity is required for physiological Wnt signaling. J Biol Chem 287:34167–78.

44. Glaeser K, Urban M, Fenech E, Voloshanenko O, Kranz D, Lari F, Christianson JC, Boutros M. 2018. ERAD-dependent control of the Wnt secretory factor Evi. EMBO J 37:e97311.

45. Willert K, Nusse R. 2012. Wnt proteins. Cold Spring Harb Perspect Biol 4:a007864.

46. Kispert A, Vainio S, McMahon AP. 1998. Wnt-4 is a mesenchymal signal for epithelial transformation of metanephric mesenchyme in the developing kidney. Development 125:4225–34.

47. Galli LM, Santana F, Apollon C, Szabo LA, Ngo K, Burrus LW. 2018. Direct visualization of the Wntless-induced redistribution of WNT1 in developing chick embryos. Dev Biol 439:53–64.

48. Rawadi G, Vayssière B, Dunn F, Baron R, Roman-Roman S. 2003. BMP-2 controls alkaline phosphatase expression and osteoblast mineralization by a Wnt autocrine loop. J Bone Miner Res 18:1842–53.

49. Ching W, Hang HC, Nusse R. 2008. Lipid-independent secretion of a Drosophila Wnt protein. J Biol Chem 283:17092–8.

50. Richards MH, Seaton MS, Wallace J, Al-Harthi L. 2014. Porcupine is not required for the production of the majority of Wnts from primary human astrocytes and CD8+ T cells. PLoS One 9:e92159.

51. Galli LM, Zebarjadi N, Li L, Lingappa VR, Burrus LW. 2016. Divergent effects of Porcupine and Wntless on WNT1 trafficking, secretion, and signaling. Exp Cell Res 347:171–83.

52. Chen Q, Takada R, Takada S. 2012. Loss of Porcupine impairs convergent extension during gastrulation in zebrafish. J Cell Sci 125:2224–34.

53. Kurita Y, Ohki T, Soejima E, Yuan X, Kakino S, Wada N, Hashinaga T, Nakayama H, Tani J, Tajiri Y, Hiromatsu Y, Yamada K, Nomura M. 2019. A High-Fat/High-Sucrose Diet Induces WNT4 Expression in Mouse Pancreatic β-cells. Kurume Med J 1–8.

54. Yamamoto H, Awada C, Hanaki H, Sakane H, Tsujimoto I, Takahashi Y, Takao T, Kikuchi A. 2013. The apical and basolateral secretion of Wnt11 and Wnt3a in polarized epithelial cells is regulated by different mechanisms. J Cell Sci 126:2931–43.

55. Lyons JP, Mueller UW, Ji H, Everett C, Fang X, Hsieh J-C, Barth AM, McCrea PD. 2004. Wnt-4 activates the canonical beta-catenin-mediated Wnt pathway and binds Frizzled-6 CRD: functional implications of Wnt/beta-catenin activity in kidney epithelial cells. Exp Cell Res 298:369–87.

56. Jordan BK, Shen JH-C, Olaso R, Ingraham HA, Vilain E. 2003. Wnt4 overexpression disrupts normal testicular vasculature and inhibits testosterone synthesis by repressing steroidogenic factor 1/beta-catenin synergy. Proc Natl Acad Sci U S A 100:10866–71.

57. Bernard P, Fleming A, Lacombe A, Harley VR, Vilain E. 2008. Wnt4 inhibits beta-catenin/TCF signalling by redirecting beta-catenin to the cell membrane. Biol Cell 100:167–77.

58. Louis I, Heinonen KM, Chagraoui J, Vainio S, Sauvageau G, Perreault C. 2008. The signaling protein Wnt4 enhances thymopoiesis and expands multipotent hematopoietic progenitors through beta-catenin-independent signaling. Immunity 29:57–67.

59. Kurayoshi M, Yamamoto H, Izumi S, Kikuchi A. 2007. Post-translational palmitoylation and glycosylation of Wnt-5a are necessary for its signalling. Biochem J 402:515–23.

60. Schackmann RCJ, van Amersfoort M, Haarhuis JHI, Vlug EJ, Halim VA, Roodhart JML, Vermaat JS, Voest EE, van der Groep P, van Diest PJ, Jonkers J, Derksen PWB. 2011. Cytosolic p120-catenin regulates growth of metastatic lobular carcinoma through Rock1-mediated anoikis resistance. J Clin Invest 121:3176–3188.

61. Yamamoto TM, McMellen A, Watson ZL, Aguilera J, Sikora MJ, Ferguson R, Nurmemmedov E, Thakar T, George-Lucian M, Kim H, Cittelly DM, Wilson H, Behbakht K, Bitler BG. 2018. Targeting Wnt Signaling To Overcome PARP Inhibitor Resistance. bioRxiv 378463 [Pre-print].

62. Short B. 2017. The signal hypothesis matures with age. J Cell Biol 216:1207–1207.

63. Nakamura K, Shirai T, Morishita S, Uchida S, Saeki-Miura K, Makishima F. 1999. p38 mitogen-activated protein kinase functionally contributes to chondrogenesis induced by growth/differentiation factor-5 in ATDC5 cells. Exp Cell Res 250:351–63.

64. Sikora MJ, Jabobsen BM, O’Connor DP, Riggins RB, Stires H, Oesterreich S. 2017. ILC-CORE (Collaboration, Openness, REproducibility). Open Sci Framew 19 Sept.

